# Modeling structural variations sequencing information to address missing heritability and enhance risk prediction

**DOI:** 10.1101/2025.08.07.669060

**Authors:** Xiaoxuan Xia, Junteng Wu, Zhezhi Gao, Peixiong Yuan, Sheng Ye

## Abstract

Missing heritability remains a significant challenge in genome-wide studies focused on single nucleotide polymorphisms (SNPs) when analyzing the genetic basis of complex traits. Structural variations (SVs), which span broader genomic regions and often exert larger functional impacts than SNPs, hold promise for capturing this heritability. Although graph-based pangenomes have advanced SV detection and association analysis, traditional SNP-centric frameworks typically overlook detailed SV sequence information, treating overlapping SVs as independent variants and leading to reduced statistical power. To overcome this limitation, we developed *SVrefiner*, an algorithm that aligns overlapping SVs and partitions them into non-overlapping refined SVs (rSVs) based on sequence congruence. By generating precise genotype matrices, *SVrefiner* enables more accurate association studies. When applied to human, tomato, and pig genomes, *SVrefiner* delineated 48,712 rSVs from 77,696 human SVs, 6,607 rSVs from 51,561 tomato SVs, and 5,237 rSVs from 142,784 pig SVs. Incorporating rSVs alongside SNPs, indels, and unrefined SVs enhanced heritability estimates by up to 24.47%. Further, refined analysis revealed that gene expression traits are often influenced by specific SV subregions rather than entire SVs. Notably, integration of rSVs in human datasets increased total eQTLs by 71.56%, with *cis-* and *trans-*eQTLs rising by 21.3% and 94.4%, respectively. Mean risk prediction accuracy across over 16,000 traits improved by as much as 16.8%. These findings deepen the understanding of complex trait heritability and demonstrate the utility of rSVs for genetic improvement and disease risk assessment.

## Introduction

Missing heritability, characterized by the discrepancy between heritability estimates derived from family-based studies and the proportion of variance explained by significant single nucleotide polymorphisms (SNPs) in genome-wide association studies (GWAS), highlights the limitations of SNPs in elucidating the genetic architecture of complex traits and disease risk^1–2^. SNPs, representing single-base pair (bp) mutations, capture only a fraction of genomic variation, whereas structural variations (SVs), defined as sequence alterations larger than 50 bps, are more likely to exert substantial effects on gene function and regulation^3–7^. However, current methodologies, including short-read sequencing and conventional analytical methods, fail to detect over two-thirds of SVs adequately, thereby severely underestimating their contribution to genetic variation and disease^8–11^.

Conventional read mapping against a single linear reference genome introduces reference bias, impairing accurate variant detection, especially for non-reference alleles^12^. The development of graph-based pangenome approaches that incorporate multiple genetic variants into a unified, graph-structured reference genome improves the accuracy and consistency of SV detection, particularly using short-read sequencing data^13–17^. This advancement enables more comprehensive analyses of SV-phenotype associations by leveraging existing second-generation short-read datasets^18^. Recent studies have shown that SVs identified via pangenomes not only greatly enhance heritability estimates but also play a crucial role in crop breeding for species such as tomato^19^, pig^20^, and grape^21^.

Nevertheless, existing frameworks for SV heritability estimation predominantly rely on SNP-based analytical methods that consider sequence-similar SVs as discrete, independent variants, ignoring the shared sequence content between adjacent SVs^19–21^. For example, overlapping SVs containing identical DNA segments—such as SV-3 and SV-4 in Fig. 1, which both include *AGTACAGAACCCAAG* and form a common edge in the pangenome graph—are often treated as independent variants, resulting in redundant genotype matrices and diminished statistical power in downstream analyses^22^. Such simplification masks crucial sequence-level relationships, limiting the detection of true genetic signals in association studies, heritability estimation and risk prediction.

**Fig. 1.**
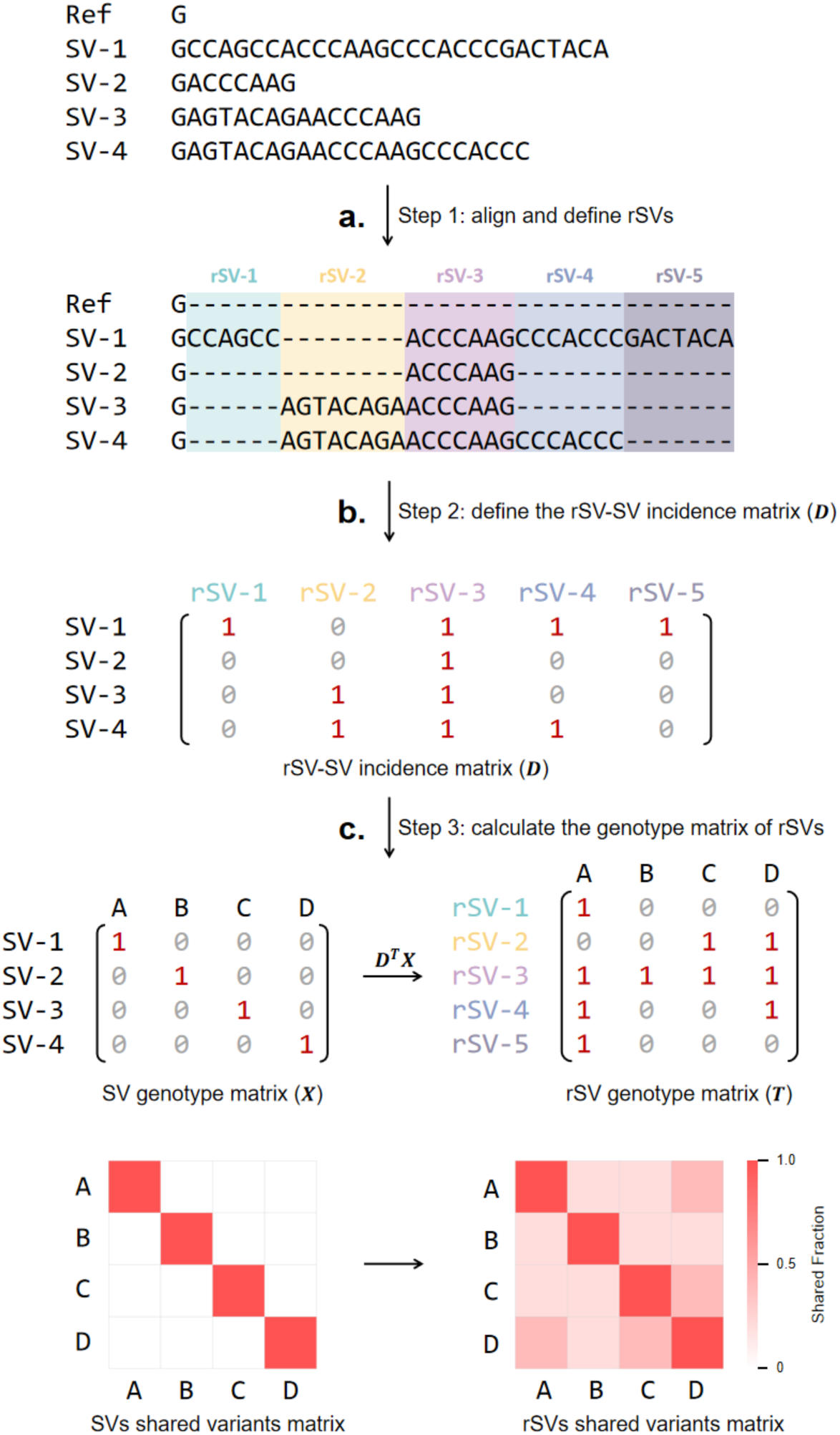
The *SVrefiner* algorithm. **a.** Step 1: Overlapped SVs are aligned to delineate rSVs. Here, SV1 to SV4 represent insertions occurring at the same genomic locus and share partially overlapping DNA sequences. By aligning these SVs, we partition their shared and unique sequences into five non-overlapping rSVs (rSV1–rSV5). **b.** Step 2: The relationships between original SVs and rSVs determined in Step 1 are encoded into an SV–rSV incidence matrix (𝑫). Matrix 𝑫 records the correspondence between each original SV and the set of rSVs. **c.** Step 3: The rSV genotype matrix (𝑻) is computed via the equation 𝑻 = 𝑫^T^𝑿, where 𝑿 represents the genotype matrix of the original SVs. The resulting rSV genotype matrix can then be analyzed using standard SNP-based analytical frameworks—such as GWAS, heritability estimation, and disease risk prediction—thus enabling more accurate downstream statistical inference.

To address this challenge, we introduce *SVrefiner*, a novel algorithm that exploits sequence-level information to decompose overlapping SVs into smaller, non-overlapping segments termed refined SVs (rSVs). This decomposition results in a refined genotype matrix, compatible with SNP-based analytical frameworks, which can more accurately capture the genetic contributions of SVs. *SVrefiner* operates through three sequential steps (Fig. 1): alignment and partitioning of adjacent SVs into unique, non-overlapping rSV intervals; construction of an incidence matrix 𝑫 representing the association between rSVs and original SVs; and generation of an rSV genotype matrix through matrix multiplication with the original SV genotype data. This approach reveals sequence-level patterns previously obscured, enabling finer discrimination of genetic backgrounds even among samples harboring distinct but related SVs.

We applied *SVrefiner* to SV datasets generated from pangenome-based references in humans, tomatoes, and pigs, each encompassing over 16,000 phenotypic traits including gene expression and metabolite profiles. Our results demonstrate that integrating rSVs with conventional genotypes markedly improves trait heritability explanation and enhances risk prediction accuracy. Additionally, rSV analysis uncovers strong genetic associations, exemplified by the gene *LINC02361*, a gene exhibiting marked differential expression between tumor and normal tissues across various cancer types, which exhibited no significant associations with SNPs, indels, or unrefined SVs, but showed a robust link with a specific rSV. Expression quantitative trait loci (eQTL) hotspot analyses further indicate that rSV signals predominantly drive association patterns in many genomic regions. These results underscore the necessity of refined modeling of SV sequences and highlight that rSV analysis should be considered an essential component in future investigations of SV–disease associations.

## Results

### High Sequence Overlap Rate in Human

Sequence overlaps among SVs are pervasive and differ markedly across species, with human SVs exhibiting the highest overlap rate at 41.50%, compared to 19.04% in tomato and 5.69% in pig. This underscores the critical need for rSV delineation to capture nuanced genomic variation. Overlapping SV clusters contain between 2 to 58 distinct variants (Fig. 2a), with sequence spans ranging from 50 bp to 116,529 bp (Fig. 2b). For example, a notable human SV cluster (Fig. 2e) on chromosome 15 (100,554,397–100,558,610 bp) containing 24 SVs (13 insertions and 11 deletions) was resolved into 46 rSVs ranging from 1 to 2,886 bp in length. Overall, the *SVrefiner* effectively partitioned 48,712 rSVs from 32,247 overlapped human SVs, 6,607 rSVs from 9,819 overlapped tomato SVs, and 5,237 rSVs from 8,130 overlapped pig SVs (Supplementary Table 1). A vast majority of human rSVs (99.76%) exhibited low linkage disequilibrium (LD; 𝑅^2^ < 0.8) with SNPs or indels, while 83.56% showed LD (𝑅^2^ < 0.8) with their original SVs (Fig. 2f). Corresponding proportions in tomato were 97.54% and 43.93% (Fig. 2g), and in pig 100.00% and 62.09%, respectively (Fig. 2h), validating that rSVs capture substantial and meaningful additional genetic information variation not sufficiently represented by conventional variant classes.

**Fig. 2.**
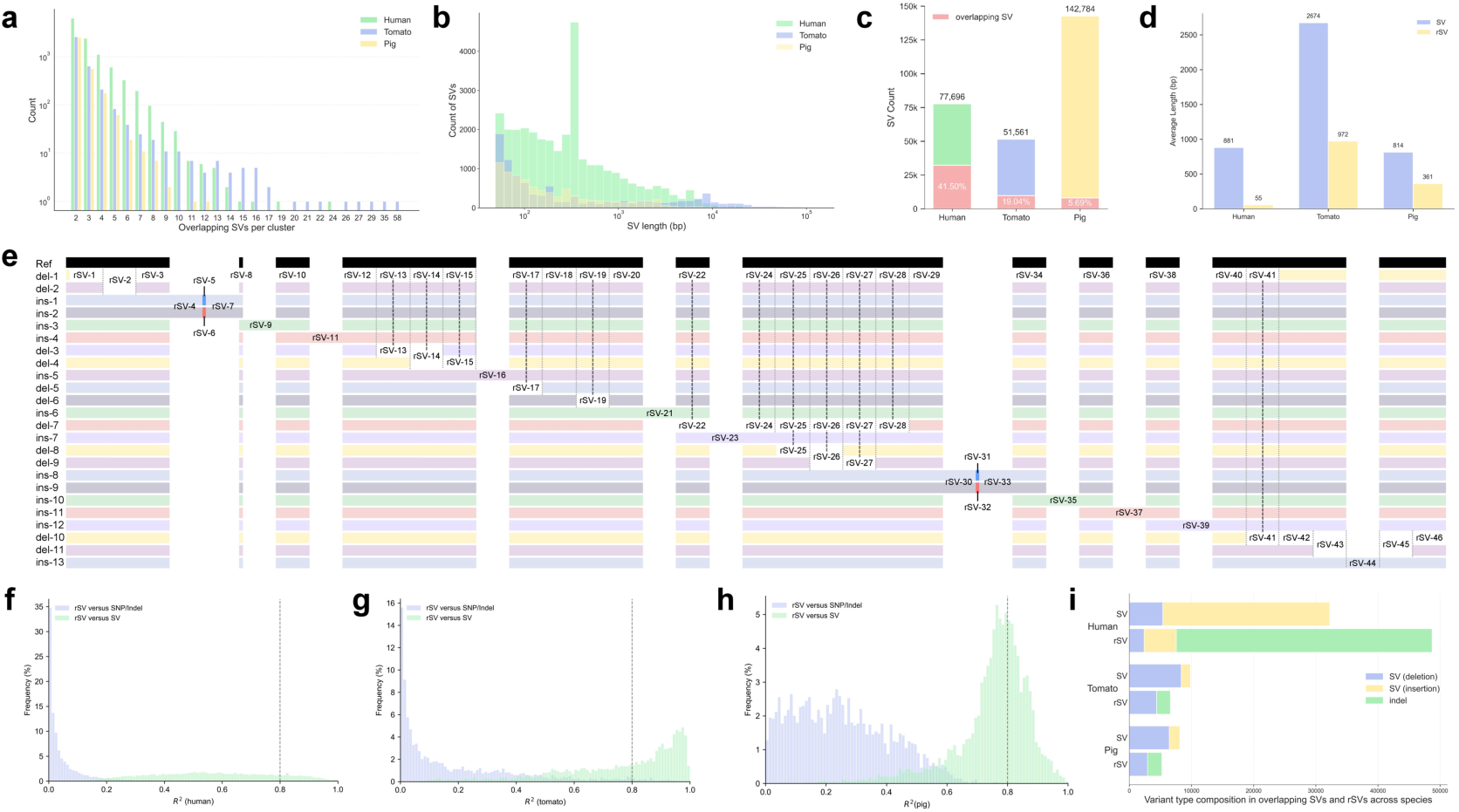
Overview of SVs and rSVs. **a.** Histogram showing the number of SVs per overlapping cluster in human, tomato, and pig. **b.** Histogram of lengths of overlapping SVs. The average length of overlapping SVs is 880.62 bp in human, 2,674.11 bp in tomato, and 813.72 bp in pig. **c.** Proportion of overlapping SVs among all SVs. **d.** Comparison of the lengths between overlapping SVs and their corresponding rSVs. The average rSV length corresponds to 6.24%, 36.35%, and 44.35% of the original SV length in human, tomato, and pig, respectively. **e.** A representative example of an SV overlapping cluster on human chromosome 15 (100,554,397– 100,558,610 bp), consisting of 24 SVs (13 insertions and 11 deletions), which were refined into 46 rSVs. Colored blocks indicate base types: A (green), T (red), C (blue), and G (orange). **f–h.** Histograms of squared Pearson correlation coefficients (𝑅^2^) between rSVs and SNPs/indels/SVs within 50 kb of the rSVs in human (**f**), tomato (**g**), and pig (**h**). For each rSV, the maximum 𝑅^2^ with adjacent SNPs/indels/SVs within 50 kb on either side is recorded^19^. **i.** Distribution of variant types before and after refinement. Most SVs (except for inversions) can be represented as combinations of insertions and deletions. We quantified the types of original SVs and rSVs in the three species, finding that the majority of SVs in human data were decomposed into indels.

### Incorporating rSVs Enhances Heritability Estimation Across Species

To assess the impact of rSVs on heritability estimation, we applied GCTA^23^ and LDAK^24^ models to 20,323 traits in tomato, 20,154 in human, and 16,323 in pig. Across both models, the inclusion of rSVs alongside SNPs, indels, and SVs significantly improved the proportion of heritable variation explained (Fig. 3a–c; Supplementary Tables 2–7). Notably, the LDAK model, which accounts for linkage disequilibrium (LD) and variant-specific characteristics, showed the greatest improvement in humans (average increase of 24.47%, 𝑃 = 1.23 × 10^-266^), with moderate yet consistent gains in tomato (3.8%, 𝑃 = 9.55 × 10^-9^) and pig (8.56%, 𝑃 = 1.19 × 10^-9^). The degree to which newly introduced variant types improve heritability estimation depends primarily on two factors: whether they are causal for the trait, and their LD with existing variants. To empirically disentangle these effects, we evaluated 100 randomly sampled overlapping SV clusters. In human and tomato, all clusters showed stable or increased heritability estimates after incorporating rSVs; in pig, this was observed in 97% of clusters, supporting the conclusion that rSVs capture additional, independent genetic signals relevant to heritability. Importantly, rSVs were found to dominate heritability for a significant subset of traits. In tomato, 17.8% of all traits were rSV-dominant (Fig. 3h), and rSV refinement led to reclassification of 41.6% of previously SV-dominant traits^19^ (Supplementary Table 8), indicating that rSVs uncover novel and functionally meaningful contributors to trait variance that are missed by traditional SV-based analyses.

**Fig. 3.**
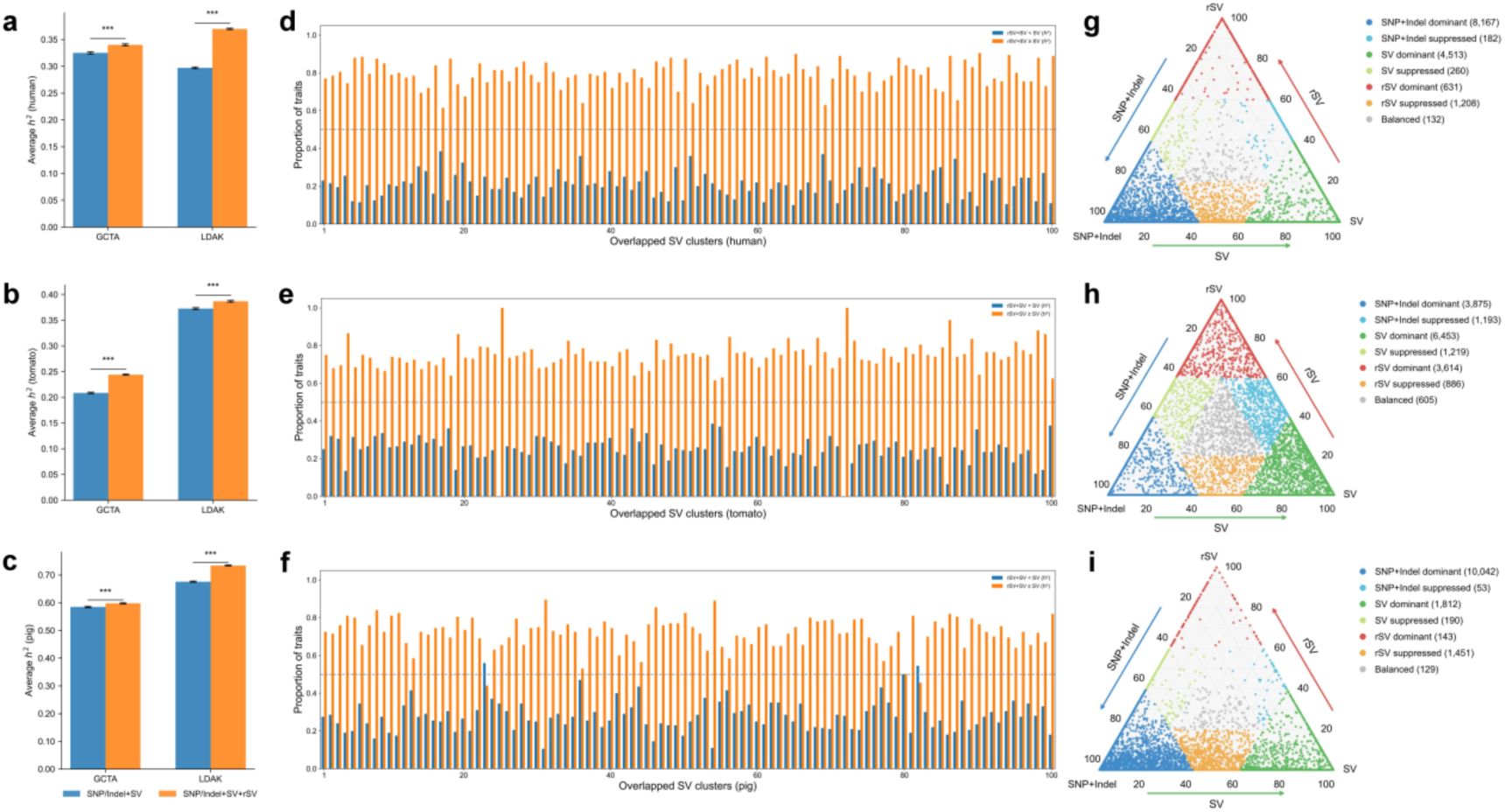
The contribution of rSVs to heritability. a, b, **c.** Comparison of heritability estimates (ℎ^2^) obtained using different combinations of genetic variants and algorithms. “SNP/Indel + SV” (blue) and “SNP/Indel + SV + rSV” (orange) refer to composite models that include either two or three categories of variants. P-values were calculated using two-sided t-tests. In humans, heritability increased by 4.66% (𝑃 = 2.10 × 10^-7^) with GCTA and by 24.47% (𝑃 = 1.23 × 10^-266^) with LDAK. In tomato, the inclusion of rSVs led to an increase in explained heritability of 17.0% (𝑃 = 2.83 × 10^-87^) and 3.8% (𝑃 = 9.55 × 10^-9^) as estimated by GCTA and LDAK, respectively. In pigs, the respective increases were 2.24% (𝑃 = 7.88 × 10^-5^) and 8.56% (𝑃 = 1.19 × 10^-86^). **d, e, f.** Comparison of the proportion of traits with higher or equal heritability when estimated using both SVs and rSVs versus using SVs alone within the same overlapping variant group. The analysis was based on 100 randomly sampled SV overlapping groups and included 200 traits under investigation. Proportion of SV clusters in which adding rSVs led to ≥50% of traits showing equal or increased heritability compared to using SVs alone. Results: 100% in human and tomato, 97% in pig. **g, h, i.** Proportion of heritability explained by different types of variants (SNPs/indels, SVs, and rSVs). A trait is defined as rSV-dominant if rSVs account for ≥60% of its total estimated heritability; analogous definitions apply to other variant types. Heritability was estimated using a composite model including SNPs/indels, SVs, and rSVs, with GCTA. Traits with an estimated heritability of ℎ^2^ = 0 (5,061 in human, 2,478 in tomato, and 2,513 in pig) are not shown. The numbers in parentheses indicate the number of traits in each group.

### rSVs Improve Association Mapping and eQTL Discovery Across Species

Association analyses revealed 12,696 rSV-trait associations in human, 272,618 in tomato, and 15,696 in pig, at stringent Bonferroni-corrected p-value thresholds. Incorporation of rSVs nearly doubled the number of significantly associated human traits (82.71% increase), with more modest though consistent gains in tomato and pig (8.92% and 10.34%, respectively), suggesting that rSV refinement reveal regulatory variants undetected by traditional SV methods. Furthermore, incorporating rSVs into existing SV datasets markedly enhanced the detection of eQTLs. Human *cis-*eQTL discovery increased by 21.3%, and *trans-*eQTLs by 94.4% with rSV integration (Fig. 4a, 4b). Similar though more moderate increases were seen for tomato and pig (Supplementary Fig. 1a, 1b, 2a, 2b). Notably, the proportion of gene expression heritability explained by *trans-*eQTLs in humans rose fivefold (0.015 to 0.081, 𝑃 = 1.02 × 10^-68^), with tomato and pig showing 16.6% (𝑃 = 1.44 × 10^-26^) and 6.3% (𝑃 = 6.78 × 10^-4^) improvements, respectively (Fig. 4c). This indicates that rSVs critically enhance functional genomic signal detection.

**Fig. 4.**
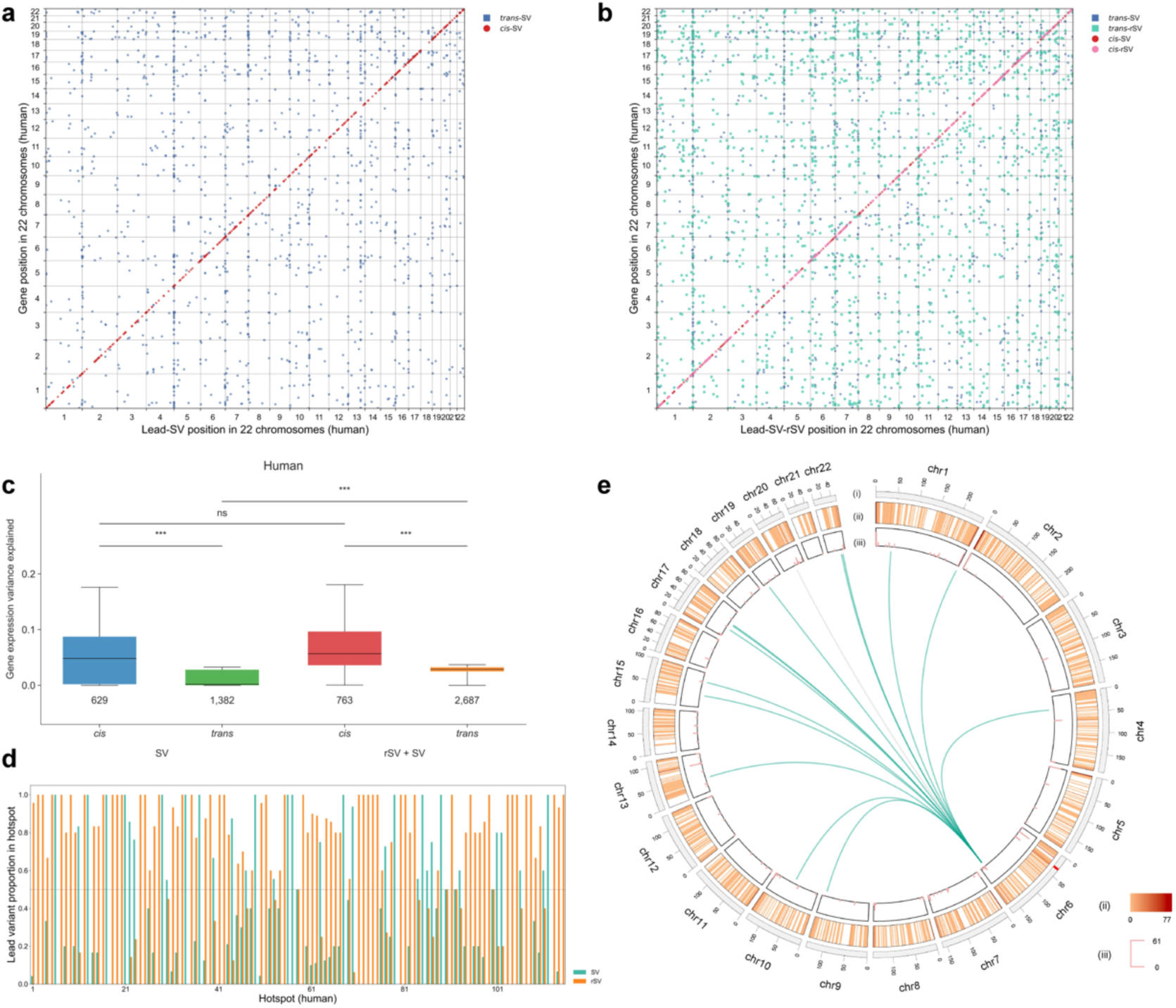
Genome-wide eQTL Identification and Hotspot Analysis in Human Chromosomes. **a**. Positions of eQTLs identified by SV. **b**. Positions of eQTLs identified by rSV and SV. **c**. Changes in the number of *cis-* and *trans-*eQTLs and their explained heritability after incorporating rSVs. The numbers below the boxplots represent the total number of detected eQTLs across all traits. The boxes represent the interquartile range (25th to 75th percentile), with the horizontal line indicating the median. Whiskers extend to the 10th and 90th percentiles. P-values were calculated using two-sided t-tests. ns: not significant; ***: 𝑃 < 0.001. Detailed results could be found in Supplementary Table 9-11. **d**. Proportion of SVs and rSVs in lead variants within all hotspots. **e**. Circle plot of 22 human chromosomes. The outermost circle shows chromosome ideograms (Mb), with the red annotation in the chr6 region indicating the MHC region. The second circle displays the number of target genes for all eQTLs in each 2-Mb window. The third circle shows the number of target genes for each *trans-*eQTL hotspot. The innermost circle highlights hotspot-31 and its target genes. Regulatory connections are shown as green lines when the lead variant is an rSV and as grey lines when it is an SV.

Hotspot analyses of *trans-*eQTL in humans identified 115 regions, with 67% being rSV-dominant hotspots (Fig. 4d, Supplementary Table 12). Gene ontology (GO) enrichment revealed that these hotspots regulate functionally related gene networks (Supplementary Table 15). For example, hotspot-31, located near the major histocompatibility complex (MHC) region, showed that rSVs account for 93% of the lead variants, and was enriched in key biological processes including gluconeogenesis, glucose metabolism, stem cell maintenance, and protein acetylation (Fig. 4e). Given the extensive LD and regulatory complexity of the MHC locus, we further investigated whether the observed *trans*-eQTL signal was primarily attributable to rSVs rather than to nearby SNPs or SVs. Fine-mapping analysis revealed that the average maximum posterior inclusion probability (PIP) was 0.004 for SNPs/indels, 0.037 for SVs, and 0.0531 for rSVs (Supplementary Table 18), indicating that rSVs are more likely to be causal variants in this region. These results support a functional role for rSVs in modulating innate immune responses. Beyond humans, the inclusion of rSVs also contributes to hotspots discovery in tomato and pig, significantly improved the heritability explained by *trans*-eQTL. Beyond humans, the inclusion of rSVs also contributed to *trans*-eQTL hotspot discovery and significantly improved the heritability explained in tomato and pig (Supplementary Note I, Supplementary Fig. 1 and 2, Supplementary Tables 13 and 14); in tomato, for instance, 7.0% of trans-eQTL hotspot lead variants were rSVs, and these hotspots were enriched in mitochondrial and oxidative stress–related pathways (Supplementary Fig. 1d, Supplementary Table 13, 16).

### rSVs Reveal Gene-expression Associations Missed by Conventional SV Analysis

Taking the human gene *ENSG00000285730* as an example, we found its expression is regulated by a cluster of overlapping insertions located in its enhancer region (Fig. 5a). While the original SVs did not show significant associations, rSV_1_4_6_9_11_13_16 (𝑃 = 2.32 × 10^#*$^) and rSV_3_8_15 (𝑃 = 1.62 × 10^#*)^), exhibited strong expression associations (Fig. 5b, c, d). Additionally, a cluster of significant SNPs/indels was identified ∼40.7 kb downstream of the transcription end site; however, this SNP/indel cluster was separated from the target gene by six intervening genes, raising concerns about its biological relevance. Fine-mapping results further supported the causal role of rSV_1_4_6_9_11_13_16 in regulating *ENSG00000285730* expression (SuSiE PIP = 0.539, PIP for rSV_3_8_15 = 0.002, Supplementary Table 19), whereas the SNP cluster yielded weaker evidence (with highest PIP = 0.022 for indel GRCh38.chr6:166961363:T:TA, Supplementary Table 19), reinforcing the utility of rSVs in pinpointing true regulatory variants.

**Fig. 5.**
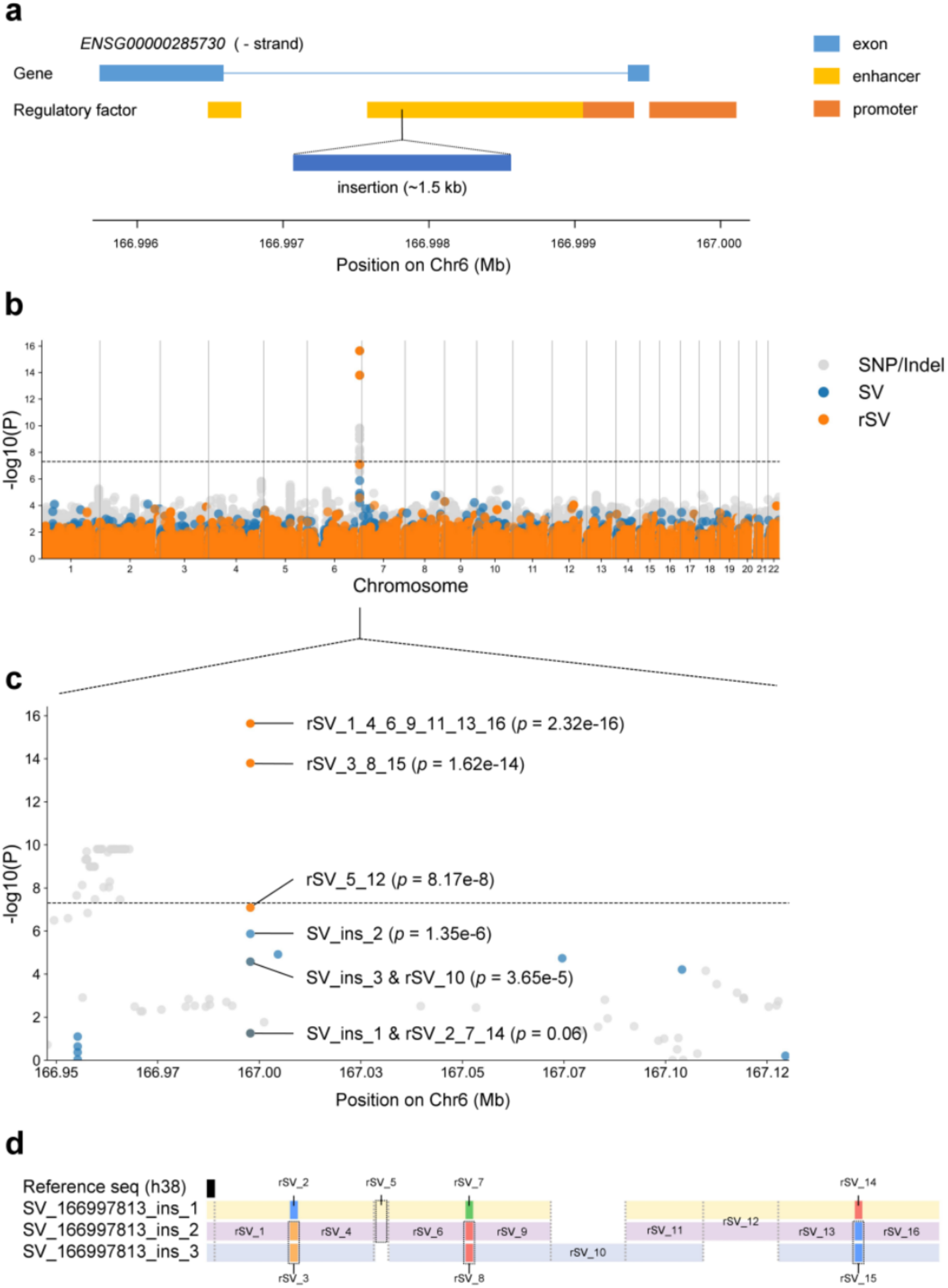
rSVs in enhancer show significant associations with gene expression in human. **a.** The gene *ENSG00000285730* is located on the negative strand of chromosome 6 in human. Blue boxes represent exons, while yellow and orange bars indicate enhancer and promoter elements, respectively. A group of insertions overlaps both the intronic region and upstream enhancers. Notably, rSVs derived from these insertions show significant association, whereas the original SVs do not, suggesting that finer-scale variants may better capture regulatory effects on gene expression. **b.** Manhattan plot of genome-wide association signals in human. Variants include SNPs/Indels (gray), original SVs (blue), and refined SVs (rSVs, orange). P values were estimated using a linear regression model adjusted for the first four principal components (PC1–PC4) to account for population stratification. The horizontal dashed line indicates the genome-wide significance threshold (𝑃 = 5 × 10^-8^, −log₁₀P ≈ 7.3). **c.** Zoomed-in view of an associated region on chromosome 6, outside of the MHC region. Multiple rSVs (e.g., rSV_1_4_6_9_11_13_16, 𝑃 = 2.32 × 10^-16^) show substantially stronger association signals compared to the original SVs. Other SVs and rSVs exhibit moderate (e.g., SV_ins_2) or weak (e.g., SV_ins_1 & rSV_2_7_14) associations. **d.** Structural decomposition of SV_166997813_ins_1 to SV_166997813_ins_3 into refined variants (rSV_1 to rSV_16). Colored blocks indicate base types: A (green), T (red), C (blue), and G (orange).

Another illustrative example is the gene *LINC02361*, which exhibits significantly differential expression in thymoma (THYM), pheochromocytoma and paraganglioma (PCPG), and kidney chromophobe (KICH). According to TCGA and GTEx data, *LINC02361* was markedly upregulated in THYM tumor tissues compared to normal samples (TPM = 11.54 vs. 0.83; Log2FC = 3.7; 𝑃 = 1.82 × 10^-1^^24^) (Fig. 6a, 6b). A similar upregulation pattern was observed in PCPG (TPM = 2.94 vs. 0.13; Log2FC = 4.5; 𝑃 = 0.011), while in KICH, the gene was significantly downregulated (TPM = 0.27 vs. 2.06; Log2FC =-2.9; 𝑃 = 3.19 × 10^-17^). The expression of *LINC02361* was strongly associated with a group of rSVs located approximately 4.6 kb downstream, among which the most significant was rSV_1_3_5 (𝑃 = 5.77 × 10^-21^), followed by rSV_4 (𝑃 = 2.99 × 10^-19^) (Fig. 6d, 6e). Although both rSVs exhibited strong statistical signals, fine-mapping analysis identified rSV_1_3_5 as the likely causal variant, with a high posterior inclusion probability (PIP = 0.792), while rSV_4 showed a much lower PIP of 0.002 (Supplementary Table 20). In addition, several SVs and SNPs also showed statistically significant associations, but their causal potential was markedly lower. The most strongly supported SV and SNP reached P-values of 4.35 × 10^-14^ (SV_54482, PIP = 0.136, Supplementary Table 20) and 3.56 × 10^-11^ (GRCh38.chr12:131940964:C:A, PIP = 0.034, Supplementary Table 20), respectively. Similar patterns were observed in tomato and pig (Supplementary Note I, Supplementary Fig. 3 and 4, Supplementary Tables 21 and 22), further highlighting the unique value of rSVs in identifying biologically meaningful regulatory variants, particularly in scenarios where traditional variant classes fail to adequately explain regulatory effects.

**Fig. 6.**
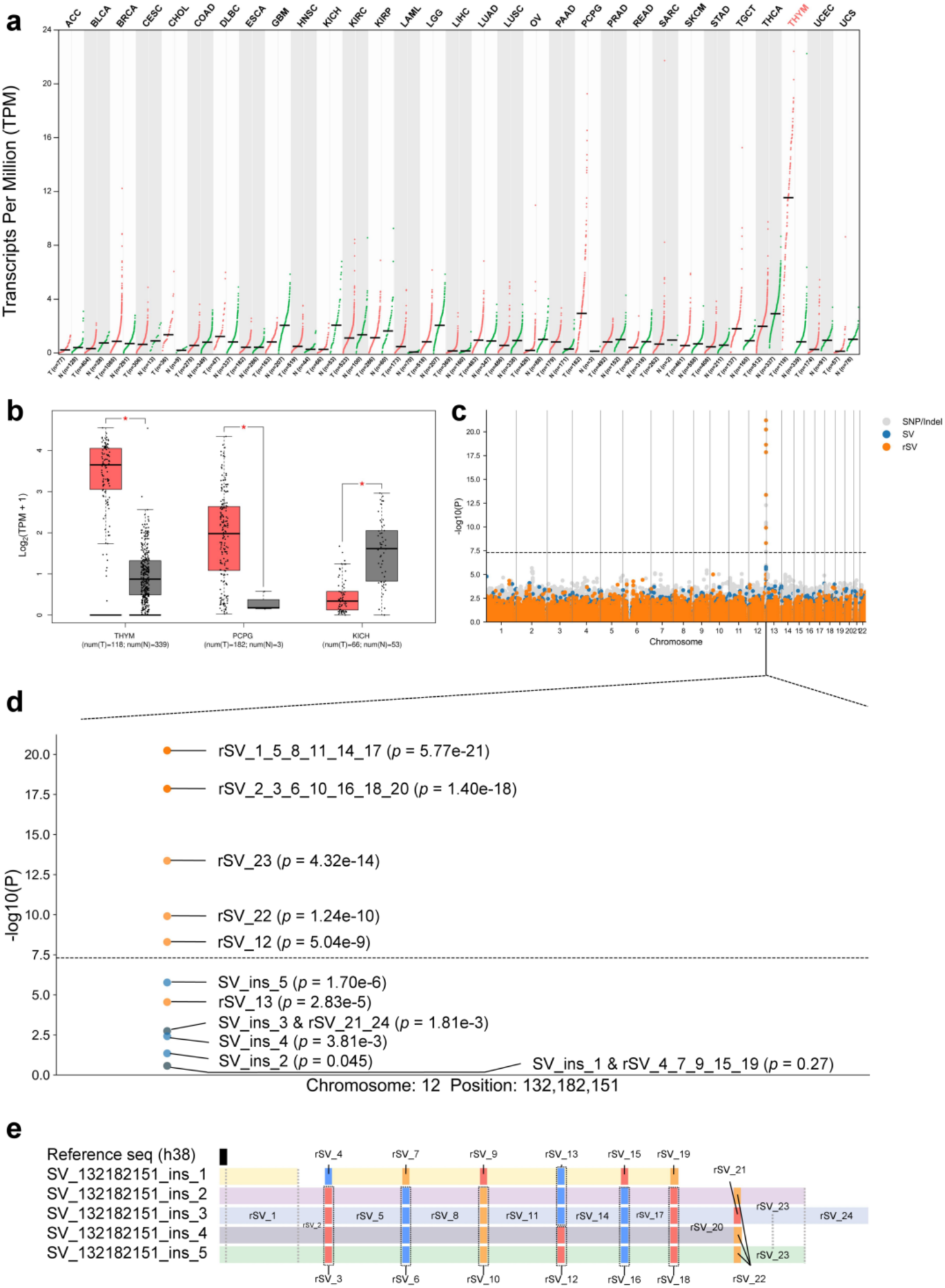
Expression of *LINC02361* across various cancer types and its association with rSVs. **a.** Expression levels of *LINC02361* across multiple cancer types in TCGA, based on TCGA and GTEx matched normal tissue data. **b.** Boxplot comparing expression levels of *LINC02361* between THYM, PCPG, and KICH tumor samples and normal tissues. Expression values were Log2(TPM+1) transformed. The gene shows a Log2FC change of 3.7 in THYM (𝑃 = 1.82 × 10^-124^), 4.5 in PCPG (𝑃 = 0.011), and-2.9 in KICH (𝑃 = 3.19 × 10^-17^). Expression differences between tumor and normal tissues were considered significant if the Log2FC was ≥ 1 or ≤ −1, with a 𝑃 < 0.05^-25^. **c.** Manhattan plot of genome-wide association analysis between genetic variants and *LINC02361* expression. A significant association peak was detected on chromosome 12. **d.** Zoomed-in view of genetic variants located at Chr12:132,182,144. Strong associations were observed for rSV_1_3_5 (𝑃 = 5.77 × 10^-21^) **e.** Alignment of three insertion SVs at Chr12:132,182,144 and their corresponding refined rSV groups. All rSVs are located approximately 4.6 kb downstream of the target gene.

### Incorporating rSVs Empowers Risk Prediction

To assess the contribution of rSVs to complex trait prediction, we computed polygenic risk scores (PRS) for 200 traits across human, tomato, and pig, using both Clumping and Thresholding (C+T) and Elastic Net models. In all three species, incorporating rSVs into models that already included covariates, SNPs, indels, and SVs consistently improved prediction accuracy. Under the C+T method, rSV inclusion led to mean R² gains of 14.3% (human), 16.8% (tomato), and 12.5% (pig); corresponding gains with Elastic Net were 11.3%, 15.3%, and 11.7% (Fig. 7a, Supplementary Tables 22–27). We further quantified variant-type contributions by comparing the mean absolute PRS effect sizes. In the C+T model, rSVs accounted for 25.78%, 38.76%, and 20.60% of the total effect in human, tomato, and pig, respectively (Fig. 7b); in Elastic Net, their contributions rose to 32.81%, 27.55%, and 25.23% (Supplementary Fig. 5). These results highlight the unique value of rSVs in capturing complementary genetic signals beyond conventional variants, thereby enhancing complex trait prediction across species.

**Fig. 7.**
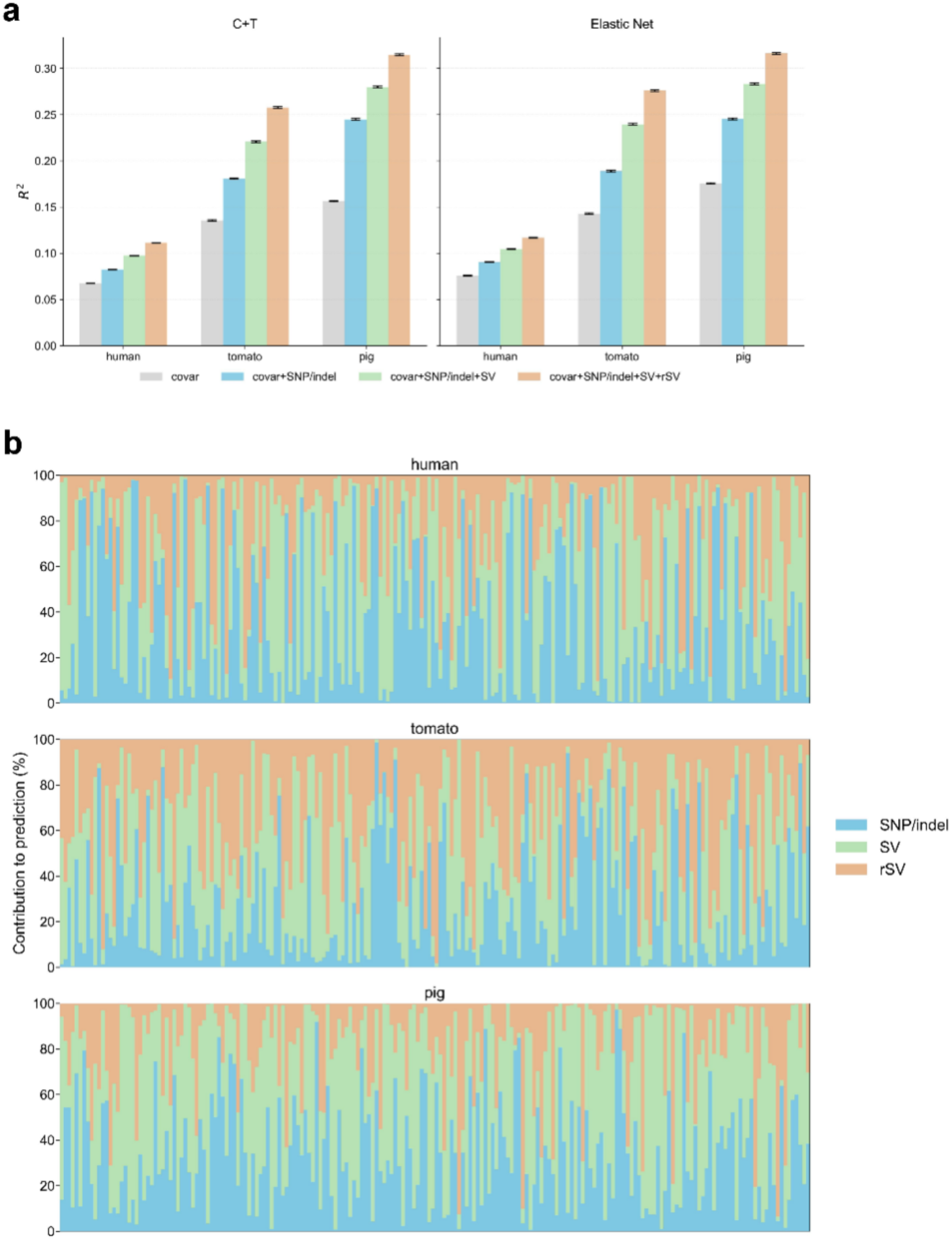
Prediction performance and variant contribution of SNP/indels, SVs, and rSVs across species. **a.** Prediction accuracy (𝑅^2^) across all traits in human, tomato, and pig using two approaches: Clumping and Thresholding (C+T, left) and Elastic Net (right), evaluated under four models. Model 1: covariates only; Model 2: covariates + PRS of SNP/indels; Model 3: covariates + PRS of SNP/indels + PRS of SVs; Model 4: covariates + PRS of SNP/indels + PRS of SVs + PRS of rSVs (see Methods for details). Bars represent the mean 𝑅^2^ across traits, with error bars indicating the standard error of the mean. **b.** Under the C+T approach (Model 4), 200 traits were randomly selected to evaluate the relative contribution of three variant types (SNP/indels, SVs, and rSVs) to trait prediction. Each stacked bar represents one trait, and the contribution of each variant type was estimated based on the absolute effect size of its PRS in Model 4. Corresponding results for the Elastic Net approach are shown in Supplementary Fig. 5.

## Computation

In aligning overlapping SVs, real data show that variant lengths range from 50 bp to 116,529 bp (Fig. 2b), and each alignment group contains 2 to 58 SVs, presenting a highly challenging multi-sequence alignment problem. *SVrefiner* addresses this by anchoring to a linear reference genome (see Methods), thereby bypassing most of the computationally intensive alignment steps. Under our tested configurations, *SVrefiner* consistently emerged as the fastest method, particularly when the maximum SV sequence length 𝐿 ≥ 10^4^ bp (Fig. 8a, b). Specifically, when 𝐿 = 10^5^ bp, *SVrefiner* achieves a 28-fold speed improvement over MAFFT^26^, a leading approximate alignment tool, and significantly outperforms both MUSCLE^27^ and Clustal Omega^28^. Furthermore, *SVrefiner* maintains consistently low memory usage across varying sequence lengths and group sizes (Fig. 8c, d). Its computational footprint remains minimal, with most resource consumption attributed to I/O operations rather than memory-intensive processes, making it highly scalable and efficient for large-scale SV datasets.

**Fig. 8.**
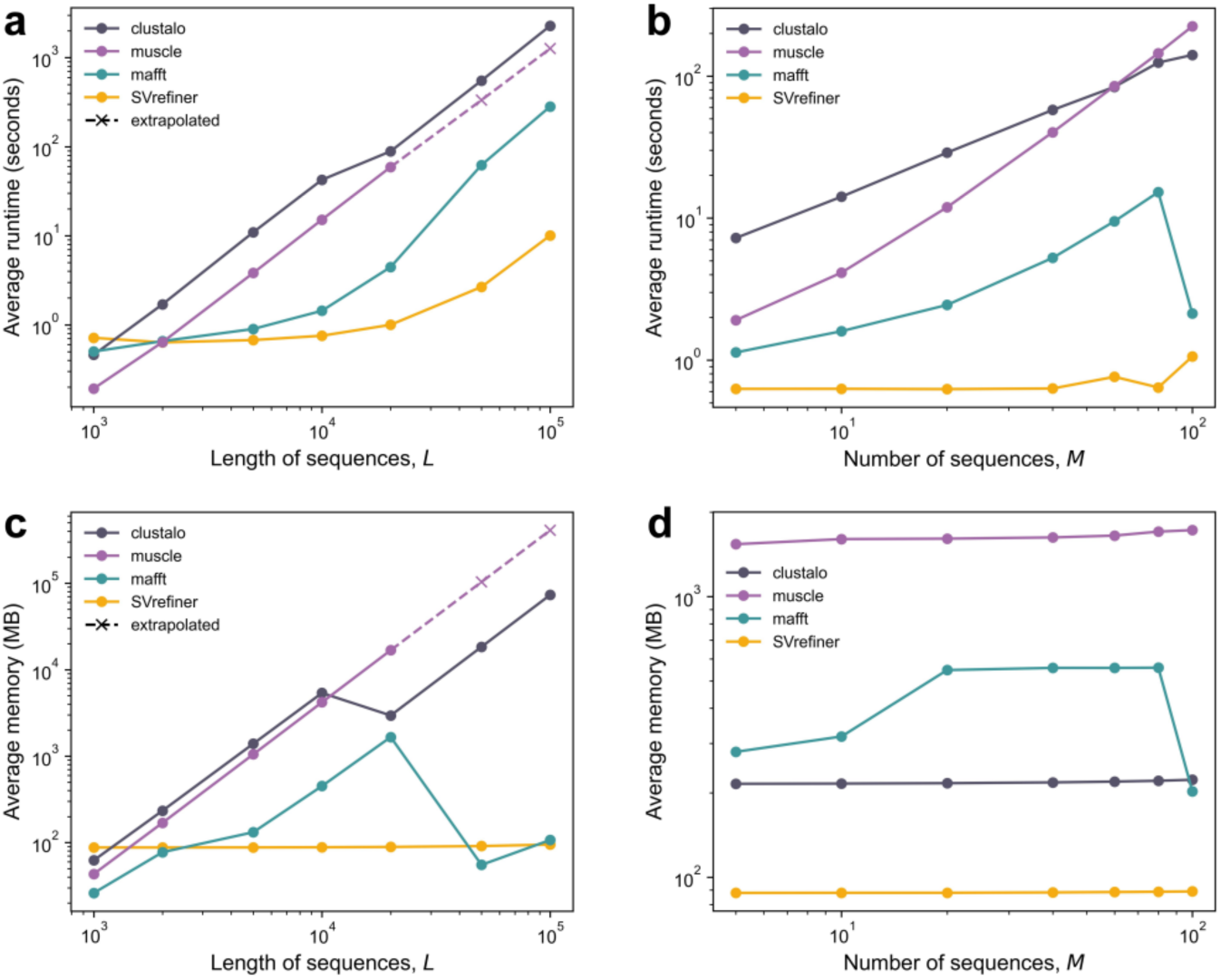
Computation time comparison. Comparison of runtime and memory usage of *SVrefiner* and three commonly used multiple sequence alignment (MSA) tools (MAFFT, MUSCLE, and Clustal Omega). Each point represents the mean of 10 independent replicates. Extrapolated points indicate values estimated by trend extrapolation due to memory limitations of the original tools. The group size (𝑀) consists of a fixed 2,000 bp reference sequence and 𝑀 50 bp deletion SVs. **a.** Average runtime as a function of sequence length (𝐿, from 1,000 to 100,000 bp) at fixed group size of 𝑀 = 2. **b.** Average runtime as a function of group size (𝑀, from 5 to 100) at fixed length size of 𝐿 = 2,000 bp. **c.** Average memory usage as a function of sequence length (𝐿) at fixed group size of 𝑀 = 2. **d.** Average memory usage as a function of group size (M) at fixed length size of 𝐿 = 2,000 bp.

## Discussion

This work introduces *SVrefiner*, an innovative algorithm that substantially advances the fine-scale modeling of SVs by integrating sequence overlap information into variant partitioning and genotype matrix construction. Traditional SV analyses, often anchored in SNP-centric frameworks, overlook the complex sequence relationships among overlapping SVs, leading to underpowered heritability estimates, diminished GWAS sensitivity, and suboptimal risk prediction. *SVrefiner* addresses these limitations by decomposing overlapping SVs into refined, non-overlapping rSVs, capturing nuanced sequence-level variation and enabling enhanced genetic analyses when integrated with conventional SNPs, indels, and SVs datasets.

Our application of *SVrefiner* across human, tomato, and pig genomes—each covering over 16,000 traits—demonstrated consistent improvements in heritability estimation and association power, validating that rSVs capture crucial genetic information masked in conventional SV matrices. This refined modeling reflects a more precise representation of genomic variation, analogous yet more complex than SNPs, which are inherently discrete variants defined by unique position and allele.

In conventional analyses of genetic variation, SNPs are typically defined by a unique combination of genomic position and alternate allele (ALT). That is, any difference in position or ALT is sufficient to categorize two SNPs as entirely distinct variants. Consequently, existing analytical frameworks treat each column in a genotype matrix as an independent variable. However, SVs are inherently more complex than SNPs. Consider, for example, two insertions that occur at the same genomic location but share partially overlapping ALT sequences: INS-1, ALT: *ATTTC, CCGGG,…, TCCCG, GTAAC, G* and INS-2, ALT: *ATTTC, CCGGG,…, TCCCG, GGAAA, GCCCC, C*. The first 50 base pairs of both variants are identical, accounting for 89.3% of the total length of INS-1. In such cases, it is clearly inappropriate to treat INS-1 and INS-2 as entirely different variants. Indeed, highly overlapping SVs are widespread across the human genome, arising from genuine biological differences, population-specific variations^29^, or limitations in SV detection methodologies.

To address this issue, one existing approach involves merging similar SVs based on sequence similarity, as implemented in tools such as Truvari^30^. These tools evaluate ALT sequence similarity and, if a predefined threshold is met, merge similar SVs into a single event. This merging strategy is reasonable in the context of reference genomes derived from short-read sequencing, where incomplete coverage of complex regions leads to gaps and imprecise SV calls. In such cases, merging partially similar SVs helps reduce noise and stabilize downstream analyses. However, with the emergence of long-read sequencing-based pangenome references, the accuracy of SV detection has significantly improved. Contemporary tools such as Pangie^31^ and Paragraph^32^ typically identify SVs by aligning short-read sequencing data from individual samples to a curated SV database obtained from long-read assemblies. The most sequence-similar SV from the database is then assigned to the sample as its representative variant. As a result, the SVs detected under this framework are considerably more accurate and less affected by detection artifacts. Given this high-resolution context, even partially similar SVs may represent biologically meaningful differences. Aggressively merging such SVs may obscure functionally relevant variation. Therefore, we advocate for a more refined, sequence-level modeling of SVs, preserving subtle differences between variants to enhance the precision of heritability estimation and trait association analysis.

Regarding whether rSVs can fully substitute traditional SVs, our statistical analyses indicate that rSVs are not yet able to serve as a complete replacement, especially in the context of heritability estimation. Across multiple species, we observed that when using a variant set composed of SNPs/indels, non-overlapping SVs (nSVs), and rSVs, genome-wide heritability estimates did not show significant improvement and in some cases were slightly reduced compared to models employing the original SNPs/indels and SVs (Supplementary Fig. 7). A likely explanation for this observation is that co-occurring rSV fragments—originating from the same original SV—carry meaningful biological information. For example, consider a 60 bp deletion SV that is decomposed into six 10 bp rSVs. While analysis of any single rSV may capture localized effects on gene expression, certain regulatory mechanisms may require the simultaneous presence of all rSV fragments to exert a phenotypic effect equivalent to that of the original SV. In such cases, the intact structural alteration represented by the original SV provides a more accurate depiction of its functional impact, whereas a fragmented rSV-based model may fail to capture this combinatorial effect fully. Therefore, although rSVs offer a higher-resolution representation of structural variation, retaining the full SV entity remains essential in certain analytical contexts, particularly when modeling the collective effects of variants on complex traits.

The current rSV analysis framework can also be effectively applied to SV summary statistics (Supplementary Note II). Specifically, if a published study has identified disease-associated SVs and provides their ALT sequence information, researchers can use the rSV quantification algorithm implemented in *SVrefiner* to decompose these SVs into their constituent rSVs. By leveraging the SV-to-rSV mapping matrix (as shown in Fig. 1c), the summary statistics associated with each SV can be projected onto the corresponding rSVs, enabling the derivation of rSV-level summary statistics. This approach allows for reanalysis of existing GWAS datasets to uncover novel rSV–trait associations without requiring access to individual-level genotype data, thus extending the analytical value of previously published SV-based studies.

This study has several limitations. First, when the number of overlapping SVs within a given cluster is small, many of the resulting rSVs tend to have identical genotype vectors across individuals, which limits their discriminative resolution. For instance, as shown in Fig. 6, rSV1, rSV3, and rSV5 cannot be distinguished from one another, as they exhibit identical genotypes across all samples. In such cases, we treat them as a composite variant (denoted rSV1_3_5), where a genotype of 1|0 indicates that all three sequence segments are simultaneously present on the first haplotype of the given individual. This limitation may reduce the precision of rSV-based association analyses in regions with sparse or highly redundant SV configurations. Second, *SVrefiner* currently does not support complex SVs, for which both the reference and alternative allele are larger than 1 bp^33^. While our cross-species analysis suggests that complex SVs occur at relatively low frequencies (e.g., 2.22% in tomato, 17.87% in humans, and 0% in pig), they can still have important biological consequences. Extending *SVrefiner* to accommodate complex SVs will be a necessary step toward comprehensive modeling of genome structure variation. The construction of pangenome references has already enhanced the accuracy and consistency of SV detection by incorporating population diversity and alternative haplotypes. Looking ahead, as the cost of long-read sequencing continues to decline, we anticipate the availability of large-scale, individual-level long-read data. This will not only allow for more accurate reference assemblies but also enable direct SV genotyping across populations. We envision a future in which SVs—and particularly their refined representations such as rSVs—will serve as a powerful complement to SNP-based studies, offering deeper insights into the genetic architecture of complex traits and enhancing the precision of disease risk prediction.

## Methods

### DNA Sequence Alignment Strategy in SVrefiner

*SVrefiner* infers rSVs directly by leveraging the positional and alternate sequence information of SVs mapped to a single reference genome, thereby bypassing most DNA sequence alignment steps. Specifically, although the SVs used in this study were detected by aligning sample sequences to a pangenome graph, a single reference genome (e.g., GRCh38 for human) can still serve as a unified coordinate system to accurately determine the locations and replacement sequences of variants. Based on the positional and length information of SVs on GRCh38, their overlaps can be directly inferred.

For overlapping SVs, we classified them into four categories according to their positions and variant types: (1) multiple deletions at the same position, (2) multiple insertions at the same position, (3) multiple deletions at different positions, and (4) multiple insertions at different positions. For groups containing only deletions (i.e., types (1) and (3)), alignment is performed by directly replacing the deleted regions with gap placeholders (Supplementary Fig. 6, Step 3: del-1 and del-2), eliminating the need for external multiple sequence alignment (MSA) tools and thus minimizing runtime. For insertions at distinct positions (i.e., type (4)), sequence alignment can similarly be completed without invoking MSA by padding the reference with placeholders at the corresponding positions and adding matching gaps in all other sequences (Supplementary Fig. 6, Step 3: ins-2 and ins-3). In cases where multiple insertions occur at the same position, local MSA is performed using MAFFT^26^ on the inserted sequences, and the aligned segment is then inserted back into the original genomic site (Supplementary Fig. 6, Step 4: ins-1 and ins-2).

This strategy is based on the premise that if a sample sequence can be aligned to a fixed position on a single reference genome, the flanking regions are consistent with the reference. Exploiting this property allows the majority of sequence alignment steps to be avoided, with MSA invoked only for ambiguous regions, thereby substantially reducing both computational burden and memory usage while preserving alignment accuracy.

### SNPs, Indels, SVs Calling

The tomato study^19^ provided the tomato pangenome (TGG1.1) and short-read sequencing data for 332 accessions, but did not report sample-level variant calls. Similarly, the pig study^20^ provided the pig pangenome reference (pig pangenome) along with short-read sequencing data for 297 accessions. Since only PAVs were included in the pig pangenome, we obtained the SNPs/indels of each accession from a published study^35^, as was done in the pig study. For humans, the data consisted of 731 individuals from the 1000 Genomes Project (1KGP)^36^, and their variant calls were directly obtained from the HPRC pangenome study^12^.

For datasets requiring variant calling from short-read sequencing data, we adopted the same variant calling strategy used in the tomato study. 1. Sequence alignment: Raw next-generation sequencing data were mapped to the pangenome reference using vg^16^ (v1.40) to best reproduce the processing pipeline of the tomato study. The resulting BAM files were sorted using SAMtools^37^ (v1.2.1). 2. SNPs/indels calling: DeepVariant^38^ (v1.0.0) was applied in pretrained mode (--model_type WGS) to detect SNPs and indels for each sample. Only variants meeting all of the following criteria were retained: (1) total sequencing depth between 810 and 3,124; (2) quality score ≥ 20; (3) biallelic variants; (4) indels with length < 50 bp. 3. Structural variant genotyping: Structural variants (SVs) were genotyped using Paragraph^32^ (v2.4a) with default parameters.

### Transcriptome Profiling and Quantification

The pig study^20^ did not provide directly usable gene expression data. To quantify gene expression level, we use the Sscorfa11.1 transcript annotations (downloaded form NCBI https://www.ncbi.nlm.nih.gov/datasets/genome/GCF_000003025.6/) and Salmon^39^ (v1.10.1) for expression quantification. We first generated a Salmon index for Sscrofa11.1 transcript GFF file using “salmon index” with the “--gencode” flag. For each of the 297 accessions, we quantified transcript-level expression on raw RNA-seq reads using “salmon quant” with default arguments. This produces, for each library, transcript-level estimates of read counts and TPM. Finally, these transcript-level estimates were summed to gene-level estimates using tximport^40^ (v1.32) in R.

Quantification of lowly expressed genes may be confounded by sequencing noise, leading to false-positive results in subsequent analyses. To avoid this issue, we only included genes with ≥6 counts and ≥0.1 TPM in at least 20% (59/297) of samples. After filtering, 16,323 expressed genes were remaining.

### Quality Control

A total number of 731 human samples from 1KGP were extracted, representing 26 globally-distributed populations across five continental groups. Similarly, 332 samples of tomato^41^ from red-fruited clade (*S. pimpinellifolium, S. lycopersicum var. cerasiforme and S. lycopersicum*) were extracted. For pig, 297 hybrid offsprings^35^ of four indigenous Chinese pig breeds (Erhualian, Bamaxiang, Tibetan, Laiwu) and four commercial European/American pig breeds (Landrace, Large White, Duroc and Piétrain) were extracted.

Quality control (QC) was performed using PLINK^42^. We first removed variants with genotype call rate < 80%, or MAF < 0.01. To reduce multicollinearity, we pruned the rSVs by (1) removing nearby variants with LD > 0.999 using parameter--indep-pairwise 100 1 0.999; (2) removing rSVs that were in perfect LD (LD = 1) with original SVs. After QC, the numbers of remaining SNPs/indels, SVs and rSVs were 1,440,336, 77,696 and 48,712 in human, 29,353,214, 142,784, and 5,237 in pig and 7,067,407, 51,561 and 6,607 in tomato, respectively. In our study, traits including 20,154 expression traits in human, 20,323 traits (19,353 expression traits and 970 metabolite traits) in tomato and 16,323 expression traits in pig were analyzed.

### Genome-wide Association Analysis

GWAS was performed with PLINK 2.0 using linear model for each variant. Principal components were calculated using SNPs and indels, and the first four principal components were included as covariates to control for population stratification. Genome-wide significance was defined as 𝑃 < 5 × 10^-8^.

### Heritability Estimation

GCTA^23^ and LDAK^24^ were used to estimate the proportion of phenotypic variance explained by genetic variants. Both models are based on the linear mixed model:

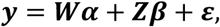

where 𝒚 is the phenotype vector, 𝑾 represents the covariates matrix, 𝒈 is the total genetic effect assumed to follow 𝑁(𝟎, 𝑲𝜎_g_^2^), and 𝜺 is the residual error, which is assumed to follow 𝑁(𝟎, 𝑰𝜎_g_^2^). The genetic relationship matrix 𝑲 summarizes the genetic similarity between individuals and was computed as:

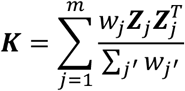

where the genotype vector 𝒁_-_ at variant 𝑗 is defined as:

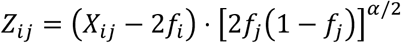

with 𝑋_ij_ being the allele count of individual 𝑖 at variant 𝑗, 𝑓_i_ the minor allele frequency (MAF) of variant 𝑗, and 𝛼 a parameter controlling the weight on variants with lower MAF. In the GCTA model, weights were set equal (𝑤_-_ = 1, 𝛼 = −1), assuming all variants contribute equally. In the LDAK model, weights were set as 𝑤_j_ ∝ 𝑢_j_, where 𝑢_j_ denote the LD weight, assigning lower weights to variants in high-LD regions. Variance components (𝜎_g_^2^ and 𝜎_e_^2^) were estimated using restricted maximum likelihood (REML) and heritability was calculated as ℎ^2^ = 𝜎_g_^2^/(𝜎_g_^2^ + 𝜎_e_^2^).

To partition heritability by variant type, kinship matrices were derived separately for SNP/indel, SV, rSV groups, and a composite model with multiple kinship matrices was fitted. For the GCTA model, variants were pruned to remove nearby high-LD variants using PLINK 2.0. For the LDAK model, the parameter’--constrain YES’ was used to prevent negative heritability estimates. The first four principal components were included as covariates to control for population stratification^19,21^.

### Fine-mapping analysis

Fine-mapping was performed with susieR^43^ (v0.14.2) using default settings. We analyzed a ±1.5 Mb window centered on the strongest significant signal. Within each group of rSVs, variants with perfect LD (LD = 1) were removed, leaving the first rSV included in analysis. For each variant, the PIP value was reported, which was used to prioritize putative causal variants.

### Polygenic Risk Prediction

Polygenic risk scores (PRS) were calculated using PLINK 2.0 and LDAK v6.1. SVs with 𝑟^2^> 0.99 to SNPs/indels and rSVs with 𝑟^2^ > 0.99 to SNPs/indels or other SVs were removed to minimize the impact of highly correlated variants. Two complementary PRS approaches were evaluated: Clumping and Thresholding (C+T) and Elastic-net shrinkage. The C+T approach can be considered a hard-thresholding method because it retains only SNPs with p-values below a specified threshold, setting the effect sizes of all other SNPs to zero. In contrast, Elastic-net is a soft-thresholding approach that constrains the total coefficient size by shrinking some SNP effect sizes toward non-zero values. Specifically, we used

1. Clumping and Thresholding (C+T) in PLINK 2.0: GWAS summary statistics were filtered using the--clump option with parameters--clump-p1 1,--clump-r2 0.2, and--clump-kb 50 to avoid overfitting caused by highly correlated variants. After clumping, variants were filtered at various p-value thresholds (0.001, 0.01, 0.05, 0.1, 0.2, 0.3, 0.5, and 0.9) using--q-score-range, and PRS were computed with the--score function.
2. Elastic-net shrinkage in LDAK v6.1: Variant effects were estimated using--elastic with two parameters to tune, 𝑝 and 𝐹, 𝑝 determines the contribution of lasso component, while 𝐹 determines the expected contribution to variance from the ridge regression component^44^. Risk scores were calculated using--calc-scores.

For both methods, samples were randomly divided into training (90%) and test (10%) sets. Within the training set, 20% of the data was further set aside as a validation set. Five-fold cross-validation was performed to select the optimal p-value thresholds in C+T and the penalty parameters in Elastic-net. The final models were trained on the entire training set and evaluated on the independent test set. Prediction performance was assessed using four nested linear models:

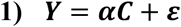

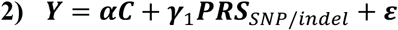

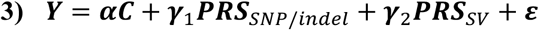

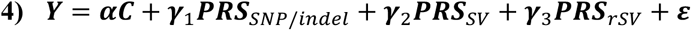

where 𝒀 denotes the phenotype vector; 𝑪 is the covariate vector containing the top four principal components, 𝑷𝑹𝑺_SNP/indel_, 𝑷𝑹𝑺_SV:_, and 𝑷𝑹𝑺_rSV_ represent the polygenic risk score vectors for SNPs/indels, SVs, and rSVs respectively; 𝜸_1_, 𝜸_2_, 𝜸_3_ are the effect sizes for each variant type, and 𝜺 denotes the residual error. For each model, we report the prediction accuracy 𝑅^2^ (defined as the squared Pearson correlation between observed and predicted phenotypes) and quantify the contribution of each variant type by the absolute value of its PRS effect size.

### eQTL Analysis

This study utilized gene expression data from three species—tomato, human, and pig—for analysis. To evaluate the changes in expression quantitative trait loci (eQTL) identification results before and after integrating rSVs into SVs, a flexible significance threshold of 1/P (where P is the total number of tested variants) was applied^45^. The first four principal components, consistent with previous analyses, were included as covariates. SVs or rSVs that were significantly associated with gene expression were grouped into candidate eQTLs if they formed a cluster of at least two significant variants with ≤50 kb between consecutive variants. Within each candidate eQTL, the variant with the smallest p-value was defined as the lead variant. Among candidate eQTLs in strong LD (𝑟^2^> 0.2), only the most significant candidate eQTL was retained to represent the cluster. If the lead variant was located within 1 Mb of the transcription start site (TSS) of the target gene, it was classified as a *cis-*eQTL; otherwise, it was considered a *trans-*eQTL. eQTL hotspots were identified using the Hotscan^46^ program (version v.05Oct2013), with a window size of 200 kb and an adjusted p-value threshold of 0.05. Gene Ontology (GO) enrichment analysis of genes regulated by *trans-*eQTL hotspots was then performed using the R package clusterProfiler^47^ (v3.10.1).

### Pan-Cancer Expression Analysis

Gene expression analysis for THYM-specific expression was conducted using the GEPIA web server^48^, which integrates data from the TCGA and GTEx datasets without requiring direct access to the raw TCGA/GTEx data. Specifically, the gene of interest and cancer subtype were selected as inputs to generate the corresponding expression profiles. Differential expression between tumor and normal tissues was evaluated using a |Log2FC| threshold of 1, indicating a significant fold change of at least two fold. A 𝑃 < 0.05 was considered statistically significant. Boxplots and expression profiles were generated to visualize the results.

### Alignment Benchmark

To evaluate the computational efficiency and scalability of *SVrefiner* in comparison with widely used multiple sequence alignment (MSA) tools, we benchmarked it against MAFFT (v7.526), MUSCLE (v5.3), and Clustal Omega (v1.2.4). All tools were installed via conda and executed with multi-threaded configurations (using the--auto option for MAFFT and Clustal Omega, and--align for MUSCLE). The benchmarks were conducted on a Linux server equipped with an Intel® Xeon® Gold 6240 CPU @ 2.60 GHz (x86_64 architecture), 128 GB of RAM, and 10 CPU cores.

To assess scalability, we generated synthetic datasets and evaluated performance across two dimensions: the number of sequences and the sequence length.

1. Sequence number benchmarks: 2,000 bp reference sequence was used to construct SV groups containing fixed 50 bp deletions. The groups included 5, 10, 20, 40, 60, 80, and 100 SV sequences.
2. Sequence length benchmarks: Two fixed sequences were simulated, each containing SV sequences with lengths of 1,000, 2,000, 5,000, 10,000, 20,000, 50,000, and 100,000 bp.

Input files were formatted as FASTA for MSA tools, and as VCF along with a reference FASTA for *SVrefiner*. For each configuration, we recorded runtime and memory usage. All benchmarks were performed with ten independent replicates to ensure statistical robustness.

## Data Availability

For tomato, the reference TGG1.1 was downloaded from the SolOmics database (http://solomics.agis.org.cn/tomato/ftp). Short-read sequencing data of 332 tomato accessions were downloaded from NCBI (BioProjects: PRJNA259308, PRJNA353161, PRJNA454805 and PRJEB5235).

For pig, the linear reference Sscrofa11.1 was downloaded from NCBI (https://www.ncbi.nlm.nih.gov/datasets/genome/GCF_000003025.6/). Small variants of 297 accessions was downloaded from the GVM (http://bigd.big.ac.cn/gvm/getProjectDetail?project=GVM000310). WGS data of these accessions was downloaded from GSA database under accession number CRA006240.

For human, the reference GRCh38.p13 was doanloaded from NCBI (https://www.ncbi.nlm.nih.gov/datasets/genome/GCF_000001405.39/). Small variants was downloaded from https://console.cloud.google.com/storage/browser/brain-genomics-public/research/cohort/1KGP/vg/graph_to_grch38/vcf?invt=AbuWsQ&pageState=(%22StorageObjectListTable%22:(%22f%22:%22%255B%255D%22)). SVs was downloaded from zenodo (https://zenodo.org/records/6797328) and processed as recommended.

The gene expression data of tomato and human was downloaded from the SolOmics database (http://solomics.agis.org.cn/tomato/ftp) and zenodo (https://zenodo.org/doi/10.5281/zenodo.10535719) respectively. Transcriptome sequencing data of pig was downloaded from GSA database under accession number CRA006240.

## Code Availability

Software for running *SVrefiner* is available at https://github.com/Lostmet/SVrefiner. All codes to reproduce the results in our paper could be found at https://github.com/Lostmet/rSV_research_pipeline.

## Acknowledgments

This work was supported by in part by Ministry of Science of Technology (2025YFC3400193 to S.Y.), the Independent Innovation Fund of Tianjin University (“Ji Jichu” Project, 2025XJ21-0003 to X.X.), the National Natural Science Foundation of China (12471275 to X.X.) and the Tianjin Natural Science Foundation (24JCQNJC01720 to X.X.). We thank Dr. Yingying Wei from the Chinese University of Hong Kong for giving precious suggestions about this project. We also thank Dr. Yao Zhou and Dr. Jian-Feng Liu for their helpful interpretations of the genomic data used in this study, and Dr. Doug Speed for his clarification and guidance regarding the use of the LDAK software.

## Author contributions

X.X. conceived and designed the study. J.W. and P. Y. wrote the software. J.W., Z. G. and P. Y. conducted data analysis. J.W. and Z. G. prepared the figures. X.X., J.W. and Z. G. co-wrote the original manuscript. S.Y. supervised the whole project. All authors read and approved the final manuscript.

