## Supplementary Notes for "Modeling structural variations sequencing information to address missing heritability and enhance risk prediction"

### Supplementary Note I

#### eQTL results in tomato

In tomato, the inclusion of rSVs led to an increase in eQTL discovery, with *cis*-eQTLs and *trans*-eQTLs rising by 2.1% and 15.9%, respectively (Supplementary Fig. 1a, 1b). Among the 574 identified *trans*-eQTL hotspots, 7.0% of the lead variants were rSVs (Supplementary Fig. 1d, Supplementary Table 13). Although rSVs account for only a small fraction of all variants, their inclusion significantly improved the explained heritability. Specifically, the heritability explained by *trans*-eQTLs increased by 16.6% ($P = 1.44 \times 10^{-26}$; Supplementary Fig. 1c), indicating that rSVs provide complementary regulatory information beyond that captured by conventional SVs. These findings suggest that rSVs may exert strong regulatory effects at specific loci and contribute disproportionately to gene expression variability. Functional enrichment showed that these rSV-associated hotspots are involved in mitochondrial and oxidative stress-related pathways (Supplementary Table 16). For example, hotspot-124, where 47% of lead variants were rSVs (Supplementary Fig. 1e), was enriched in mitochondrial ribosome assembly, genome maintenance, translation, and reactive oxygen species (ROS) metabolism—highlighting the potential of rSVs to reveal regulatory signals linked to cellular energy and stress responses.

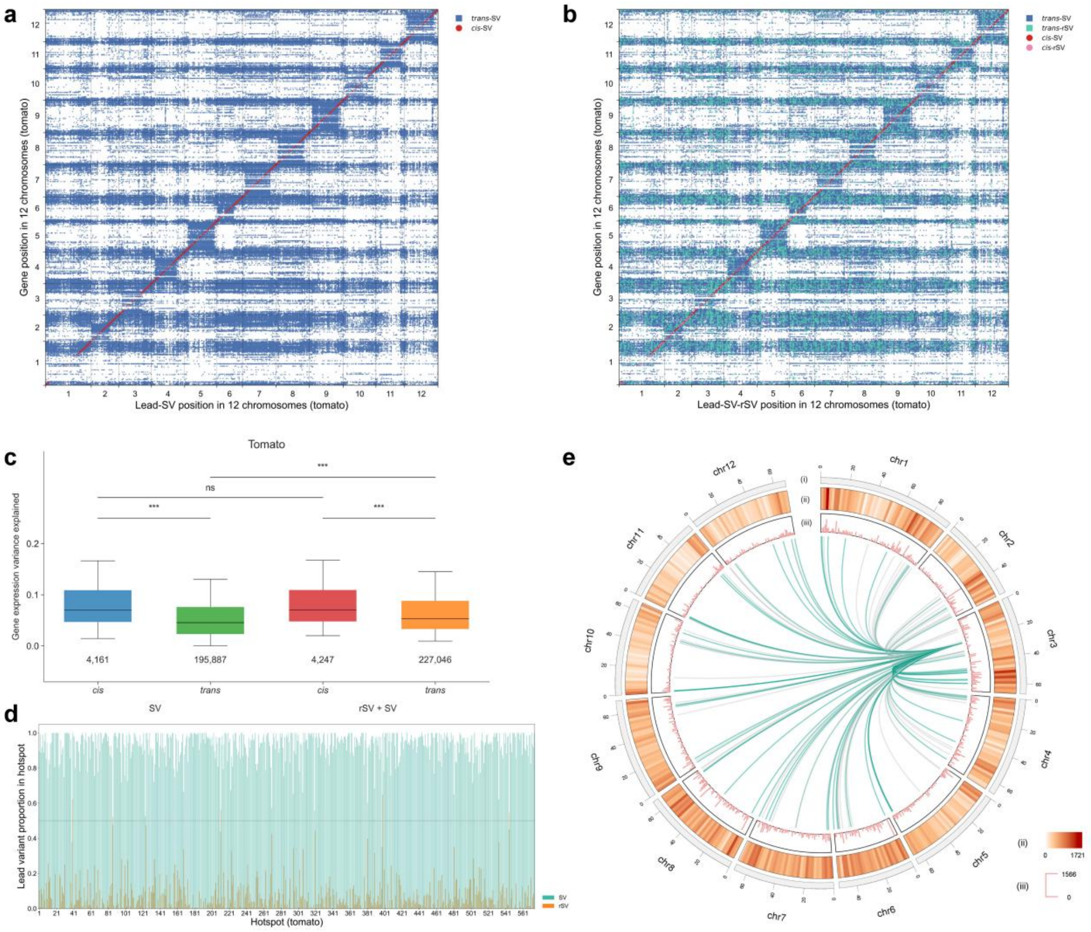


**Supplementary Fig. 1 | Genome-wide eQTL Identification and Hotspot Analysis in Tomato Chromosomes.**

**a.** Positions of eQTLs identified by SV. **b.** Positions of eQTLs identified by rSV and SV. **c.**

Changes in the number of *cis*- and *trans*-eQTLs and their explained heritability after incorporating rSVs. The numbers below the boxplots represent the total number of detected eQTLs across all traits. The boxes represent the interquartile range (25th to 75th percentile), with the horizontal line indicating the median. Whiskers extend to the 10th and 90th percentiles. P-values were calculated using two-sided t-tests. ns: not significant; ***: P-value < 0.001. Detailed results could be found in Supplementary Table 10. **d.** Proportion of SVs and rSVs in lead variants within all hotspots. **e.** Circle plot of 12 tomato chromosomes. The outermost circle shows chromosome ideograms (Mb). The second circle displays the number of target genes for all eQTLs in each 2-Mb window. The third circle shows the number of target genes for each *trans*-eQTL hotspot. The innermost circle highlights hotspot-124 and its target genes. Regulatory connections are shown as green lines when the lead variant is an rSV and as grey lines when it is an SV.

#### eQTL results in pig

In pig, the addition of rSVs resulted in a 0.38% increase in *cis*-eQTLs and a 9.6% increase in *trans*-eQTLs (Supplementary Fig. 2a, 2b), with *trans*-eQTL heritability increasing by 6.3% ($P = 6.78 \times 10^{-4}$; Supplementary Fig. 2c). Only 5.69% of original SVs showed self-overlapping sequence structures, which limited the number of rSVs that could be extracted. As a result, the improvement in heritability was smaller than in other species, but still evident. Among 215 identified *trans*-eQTL hotspots, 6.2% of the lead variants were rSVs (Supplementary Fig. 2d, Supplementary Table 14). Notably, hotspot-199 contained 19% rSVs and was enriched in genes involved in immune-related signaling, including cytokine receptor binding and intercellular communication (Supplementary Fig. 2e, Supplementary Table 17). This hotspot clustered multiple genes within coordinated immune response pathways and exhibited high connectivity among target loci, reflecting a structurally defined regulatory module. rSVs provide a complementary perspective to conventional SV annotations. When the overlap between SVs and rSVs is high, more novel sequence-level features may be recovered. However, even with low overlap—as observed in pig—rSVs can still substantially enhance the resolution of SV-related regulatory analysis.

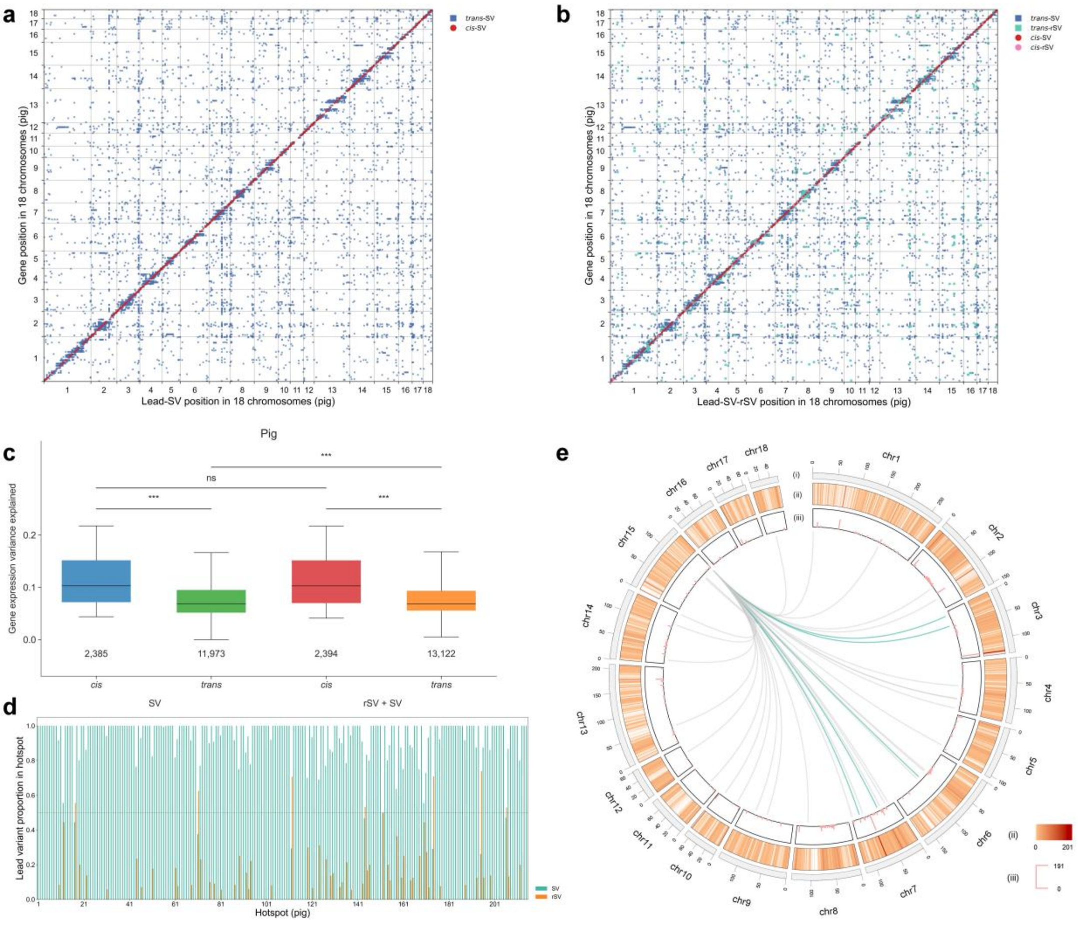


**Supplementary Fig. 2 | Genome-wide eQTL Identification and Hotspot Analysis in Pig Chromosomes.**

**a.** Positions of eQTLs identified by SV. **b.** Positions of eQTLs identified by rSV and SV. **c.** Changes in the number of *cis*- and *trans*-eQTLs and their explained heritability after incorporating rSVs. The numbers below the boxplots represent the total number of detected eQTLs across all traits. The boxes represent the interquartile range (25th to 75th percentile), with the horizontal line indicating the median. Whiskers extend to the 10th and 90th percentiles. P-values were calculated using two-sided t-tests. ns: not significant; ***: P-value < 0.001. Detailed results could be found in Supplementary Table 11. **d.** Proportion of SVs and rSVs in lead variants within all hotspots. **e.** Circle plot of 18 pig chromosomes. The outermost circle shows chromosome ideograms (Mb). The second circle displays the number of target genes for all eQTLs in each 2-Mb window. The third circle shows the number of target genes for each *trans*-eQTL hotspot. The innermost circle highlights hotspot-199 and its target genes. Regulatory connections are shown as green lines when the lead variant is an rSV and as grey lines when it is an SV.

#### rSV Regulates Expression of *Solyc01G002395* in Tomato

In tomato, GWAS results indicated that the expression of *Solyc01G002395* was associated with an rSV located on chromosome 1. No significant associations were observed for the original SVs, whereas rSV_2 showed a strong signal ($P = 1.37 \times 10^{-10}$) (Supplementary Fig. 3a–c). Fine-mapping analysis further prioritized rSV_2 as the likely causal variant, with a PIP of 0.940 (Supplementary Table 21), providing additional resolution beyond the original SV set.

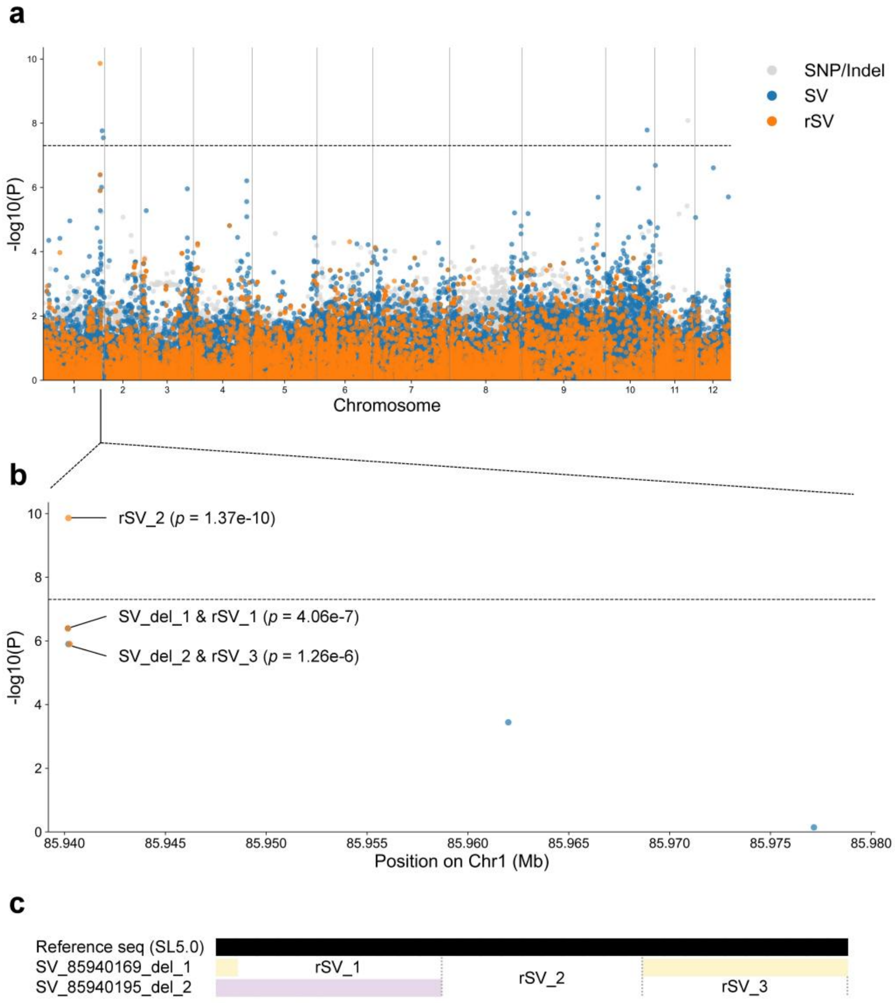


**Supplementary Fig. 3 | rSVs show stronger associations with the expression of *Solyc01G002395* in tomato.**

**a.** Association signals across all chromosomes. The P value of each variant was estimated using a linear regression model across 332 accessions. The horizontal dashed line indicates the genome-wide significance threshold ($P = 5 \times 10^{-8}$). **b.** Magnified view of the associated locus on chromosome 1, where the refined variant rSV_2 shows the strongest association ($P = 1.37 \times 10^{-10}$), while the original SVs show weaker signals. **c.** Genomic positions of SV_del_1 and SV_del_2, along with their corresponding refined variants (rSV_1 to rSV_3), are shown relative to the SL5.0 reference genome.

#### rSV Regulates Expression of *ENSSSCG00000026196* in Pig

In pig GWAS analysis, we identified a locus on chromosome 15 where the expression of *ENSSSCG00000026196* was likely associated with two rSVs. While none of the original SVs showed significant associations, rSV_2 ($P = 1.24 \times 10^{-8}$) and rSV_3 ($P = 2.38 \times 10^{-8}$) reached genome-wide significance (Supplementary Fig. 4a–c). Fine-mapping assigned the highest PIP to rSV_2 (0.341), followed by rSV_3 (0.220), whereas the top-ranking of other variants, an original SV (SV_del_2), had a PIP of 0.031 (Supplementary Table 22). These results indicate that rSVs may help refine the localization of expression-associated variants beyond what is captured by original SV annotations.

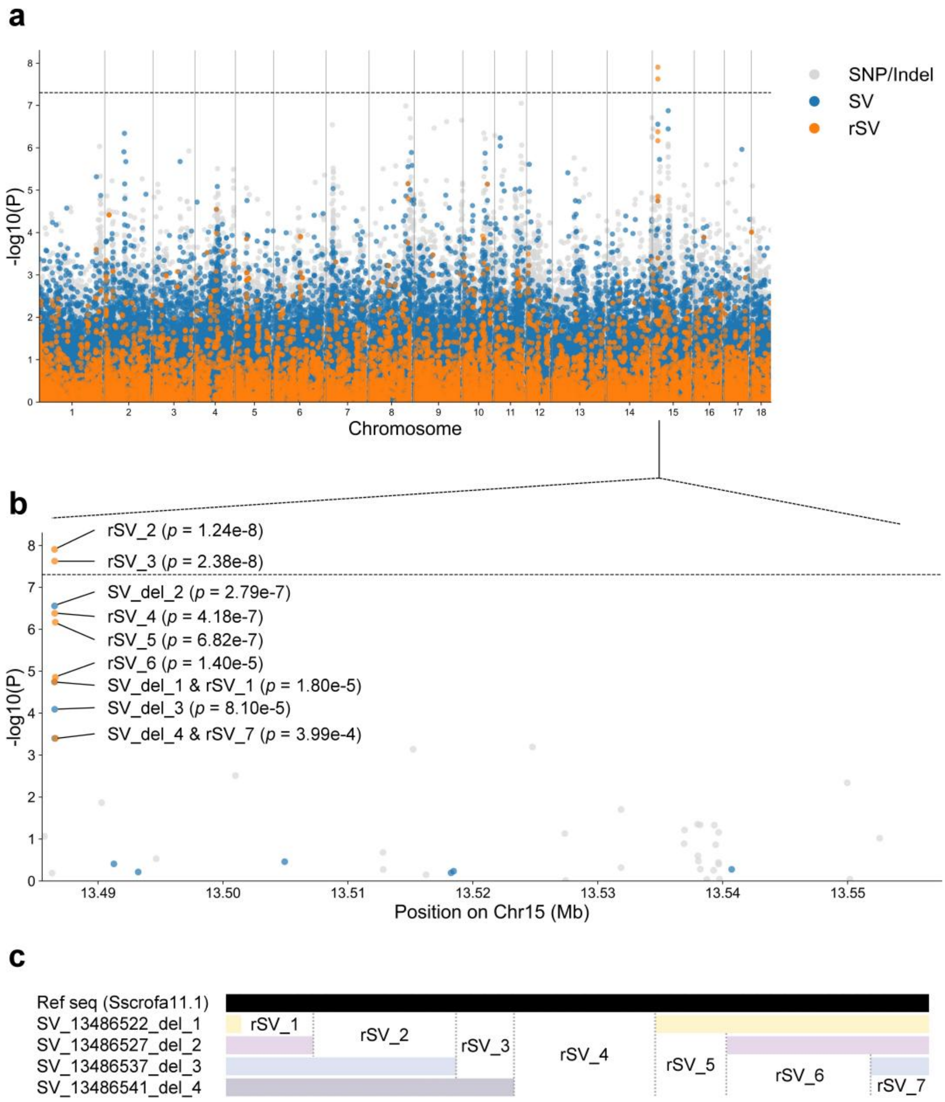


**Supplementary Fig. 4 | rSVs show stronger associations with the expression of *ENSSSCG00000026196* in pig.**

**a.** Association signals across all chromosomes. The P value of each variant was estimated using a linear regression model across 297 samples. The horizontal dashed line indicates the genome-wide significance threshold ($P = 5 \times 10^{-8}$). **b.** Magnified view of the associated locus on chromosome 15, where the refined variants rSV_2 and rSV_3 show the strongest associations ($P = 1.24 \times$

$10^{-8}$ and $P = 2.38 \times 10^{-8}$, respectively), outperforming the original SVs. **c.** Genomic positions of SV_del_1 to SV_del_4 and their corresponding refined variants (rSV_1 to rSV_7) are shown relative to the Sscrofa11.1 reference genome.

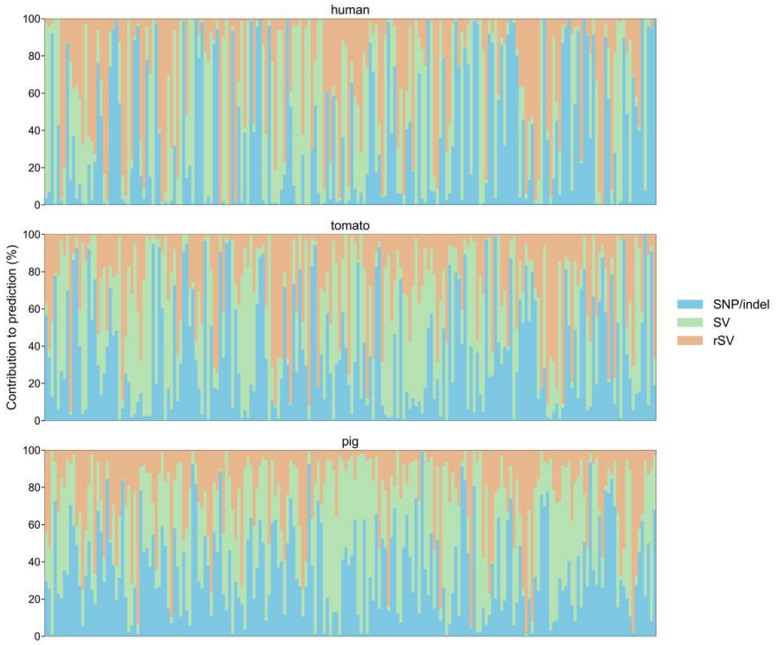


**Supplementary Fig. 5 | Variant contribution of SNP/indel, SV, and rSV across species using Elastic Net.**

Variant effect contributions were evaluated in 200 randomly selected traits using Model 4. For each trait, the relative importance of each variant type was quantified by the proportion of absolute effect sizes, as visualized in stacked bar plots. Averaged across all traits, rSVs contributed 32.81%, 27.55%, and 25.23% in human, tomato, and pig, respectively (Supplementary Tables 26–28).

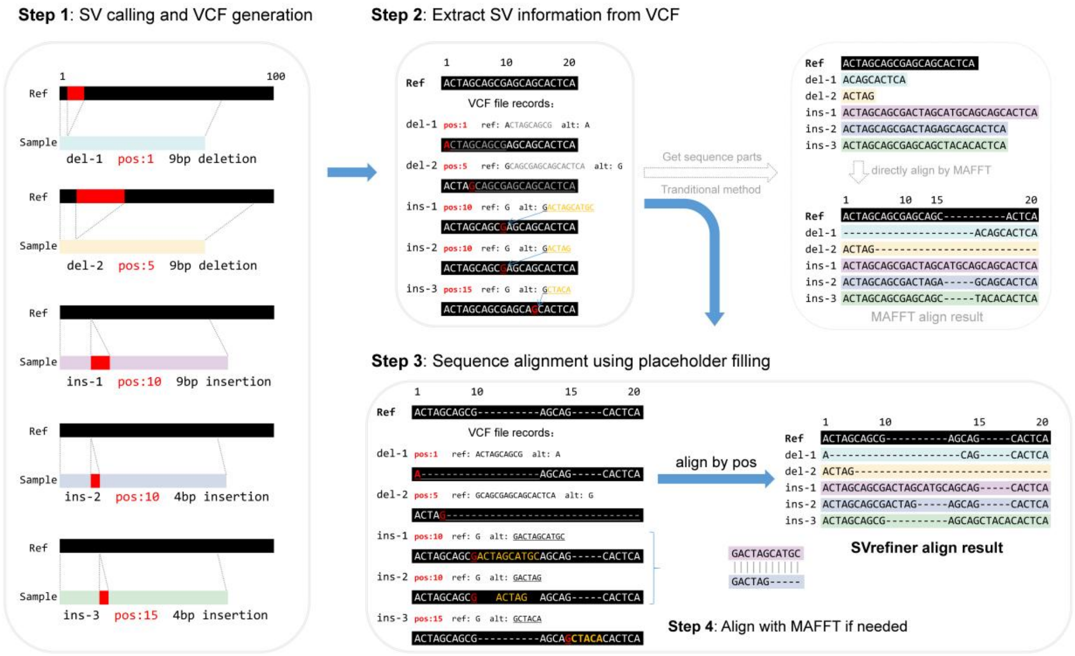


**Supplementary Fig. 6 | DNA sequence alignment scheme in *SVrefiner*.**

*Ref* represents the reference sequence. *del* denotes deletions; *ins* denotes insertions. **Step 1:** SVs are detected based on the pangenome and subsequently mapped to a single linear reference genome to determine their precise positions recorded in a VCF file. **Step 2:** SV information is extracted from the VCF and grouped by overlapping positions. **Step 3:** Sequence alignment is performed by filling placeholders after the variant positions (POS) recorded in the VCF. **Step 4:** For regions where multiple insertions occur at the same position (where POS alone cannot resolve conflicts), the segment is sliced and aligned using MAFFT, while traditional methods ignore the POS information from SV calling, leading to loss of long-fragment alignment accuracy and increased computational time.

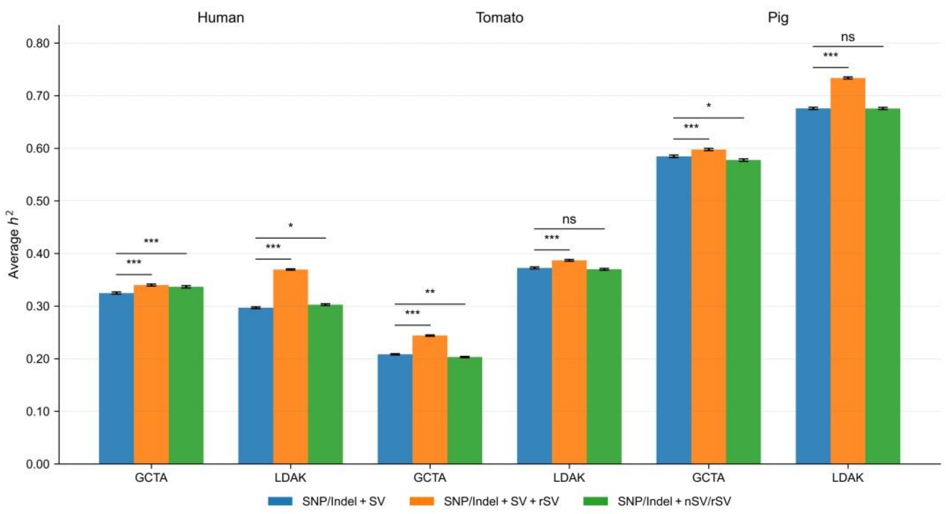


**Supplementary Fig. 7 | Comparison of heritability estimates using different variant combinations.**

We compared heritability estimates under three variant inclusion strategies: (1) SNP/Indel + SV; (2) SNP/Indel + SV + rSV; (3) SNP/Indel + nSV/rSV. Here, non-overlapping SVs (nSVs) refers to SVs that could not be decomposed into rSVs due to sequence ambiguity or the absence of shared sequence structure. Across all three species, incorporating rSVs alongside SVs significantly increased heritability estimates. In humans, adding rSVs (orange bars) resulted in a heritability increase of 4.66% ( $P = 2.10 \times 10^{-7}$ ) using GCTA and 24.47% ( $P = 1.23 \times 10^{-266}$ ) using LDAK. In tomato, the increases were 17.0% ($P = 2.83 \times 10^{-87}$) and 3.8% ($P = 9.55 \times 10^{-9}$), while in pig, they were 2.24% ($P = 7.88 \times 10^{-5}$) and 8.56% ($P = 1.19 \times 10^{-86}$), respectively. In contrast, the "SNP/Indel + nSV + rSV" model (green bars) yielded only a modest improvement in humans (up to 3.65%, $P = 6.97 \times 10^{-5}$) and even resulted in decreased heritability estimates in both tomato and pig. These findings suggest that retaining full SVs that can be decomposed and combining them with their corresponding rSVs is more effective for improving heritability estimates than relying solely on non-decomposable SVs (nSVs) and rSVs.

#### Supplementary Note II

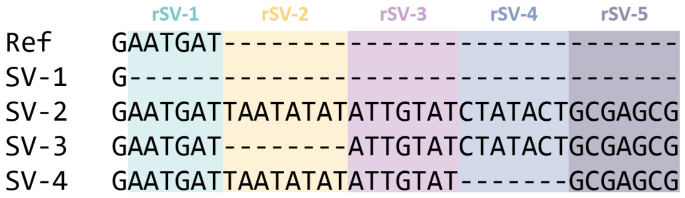


**Supplementary Fig. 8 | From SVs to rSVs.**

Suppose in this complex SV region, there are $P$ SVs, which generate $K$ rSVs. Taking Supplementary Figure 8 as an example, there are $P = 4$ SVs and $K = 5$ rSVs. This example is based on a cohort of $N = 3$ samples.

Matrix $\boldsymbol{D}_{P\times K}$:

| SV vs rSVs | $rSV1$ | $rSV2$ | $rSV3$ | $rSV4$ | $rSV5$ |
|---|---|---|---|---|---|
| $SV1$ | 1 | 0 | 0 | 0 | 0 |
| $SV2$ | 0 | 1 | 1 | 1 | 1 |
| $SV3$ | 0 | 0 | 1 | 1 | 1 |
| $SV4$ | 0 | 1 | 1 | 0 | 1 |

Matrix $\boldsymbol{X}_{N\times P}$:

| SV genotype | $SV1$ | $SV2$ | $SV3$ | $SV4$ |
|---|---|---|---|---|
| $individual1$ | 0 | 0 | 0 | 2 |
| $individual2$ | 1 | 0 | 0 | 0 |
| $individual3$ | 0 | 1 | 1 | 0 |

Matrix $\boldsymbol{T}_{N\times K}$:

$\boldsymbol{T}_{N\times K} = \boldsymbol{X}_{N\times P} \times \boldsymbol{D}_{P\times K} =$

| rSVs genotype | $rSV1$ | $rSV2$ | $rSV3$ | $rSV4$ | $rSV5$ |
|---|---|---|---|---|---|
| $individual1$ | 0 | 2 | 2 | 0 | 2 |
| $individual2$ | 1 | 0 | 0 | 0 | 0 |
| $individual3$ | 0 | 1 | 2 | 2 | 2 |

When analyzing $K$ rSVs, there are two approaches:

1. through multivariate linear regression: $y_i = \boldsymbol{t}_i\boldsymbol{\gamma} + \varepsilon_i = \sum_{k=1}^{K} t_{i,k}\gamma_k + \varepsilon_i$;
2. through single variate linear regression: $y_i = t_{i,k}\eta_k + e_i$;

Similarly, when analyzing $P$ SVs, there are two approaches:

1. through multivariate linear regression: $y_i = \boldsymbol{x}_i\boldsymbol{\beta} + \tau_i = \sum_{p=1}^{P} x_{i,p}\beta_p + \tau_i$;
2. through single variate linear regression: $y_i = x_{i,p}\theta_p + t_i$;

where $y_i$ is the phenotype of the $i$-th sample, $t_{i,k}$ is the genotype value of the $k$-th rSV in the $i$-th sample, $x_{i,p}$ is the genotype value of the $p$-th SV in the $i$-th sample, $\boldsymbol{t}_i = (t_{i,1}, \ldots, t_{i,K})$ and $\boldsymbol{x}_i = (x_{i,1}, \ldots, x_{i,P})$ are the genotype vectors, $\boldsymbol{\gamma} = (\gamma_1, \ldots, \gamma_K)$ and $\boldsymbol{\beta} = (\beta_1, \ldots, \beta_P)$ are the corresponding effect size vectors for rSVs and SVs, $\eta_k$ and $\theta_p$ are the single-variant effect sizes, and $\varepsilon_i$, $e_i$, $\tau_i$, $t_i$ are the residual errors.

Usually, in published GWAS paper, we could obtain $\hat{\theta}_p$.
Our aim is to calculate $\hat{\eta}_k$ based on $\hat{\theta}_p$ and $\boldsymbol{X}$.

We know that the estimated effect size of SVs $\hat{\boldsymbol{\beta}}$ are calculated by

$$\hat{\boldsymbol{\beta}} = (\boldsymbol{X}^T\boldsymbol{X})^{-1}\boldsymbol{X}^T\boldsymbol{Y}.$$

Therefore, the estimated effect size of rSVs $\hat{\boldsymbol{\gamma}}$ could be calculated based on $\hat{\boldsymbol{\beta}}$:

$$\begin{aligned} \hat{\boldsymbol{\gamma}} &= (\boldsymbol{T}^T\boldsymbol{T})^{-1}\boldsymbol{T}^T\boldsymbol{Y} \\ &= (\boldsymbol{D}^T\boldsymbol{X}^T\boldsymbol{X}\boldsymbol{D})^{-1}\boldsymbol{D}^T\boldsymbol{X}^T\boldsymbol{Y} \\ &= \boldsymbol{D}^{-1}(\boldsymbol{X}^T\boldsymbol{X})^{-1}\boldsymbol{X}^T\boldsymbol{Y} \\ &= \boldsymbol{D}^{-1}\hat{\boldsymbol{\beta}}. \end{aligned} \tag{1}$$

In summary statistics, we usually have marginal estimated effect size of SVs. Therefore, to obtain $\hat{\boldsymbol{\gamma}}$, we need to calculate $\hat{\boldsymbol{\beta}}$ through marginal estimated effect size $\hat{\theta}_1, \hat{\theta}_2, ..., \hat{\theta}_P$.

$$\hat{\theta}_p = \frac{\boldsymbol{x}_p^T\boldsymbol{Y}}{\boldsymbol{x}_p^T\boldsymbol{x}_p}, p = 1, ..., P.$$

Therefore,

$$\boldsymbol{x}_p^T\boldsymbol{Y} = \hat{\theta}_p\boldsymbol{x}_p^T\boldsymbol{x}_p, p = 1, ..., P.$$

From published paper, we could obtain $\hat{\theta}_p, p = 1, ..., P$. We need to calculate $\hat{\boldsymbol{\beta}}$ from $\hat{\theta}_p$ and $\boldsymbol{X}^T\boldsymbol{X}$.

$$
\begin{aligned}
\hat{\boldsymbol{\beta}} &= (\boldsymbol{X}^T\boldsymbol{X})^{-1}\boldsymbol{X}^T\boldsymbol{Y} \\
&= (\boldsymbol{X}^T\boldsymbol{X})^{-1}\begin{pmatrix} \boldsymbol{x}_1^T\boldsymbol{Y} \\ \boldsymbol{x}_2^T\boldsymbol{Y} \\ \cdots \\ \boldsymbol{x}_P^T\boldsymbol{Y} \end{pmatrix} \\
&= (\boldsymbol{X}^T\boldsymbol{X})^{-1}\begin{pmatrix} \hat{\theta}_1\boldsymbol{x}_1^T\boldsymbol{x}_1 \\ \hat{\theta}_2\boldsymbol{x}_2^T\boldsymbol{x}_2 \\ \cdots \\ \hat{\theta}_P\boldsymbol{x}_P^T\boldsymbol{x}_P \end{pmatrix}
\end{aligned} \tag{2}
$$

If when calculating $\hat{\theta}_p, p = 1, ..., P$, $\boldsymbol{x}_1, ..., \boldsymbol{x}_P$ are standardized to mean 0 and variance 1, then $\boldsymbol{x}_1^T\boldsymbol{x}_1 = ... = \boldsymbol{x}_P^T\boldsymbol{x}_P = N - 1$.

Since we have genotype data $\boldsymbol{X}$ in 1000 genome project, $\hat{\boldsymbol{\beta}}$ could be calculated by above equation (2).

Therefore, according to equation (1), the estimated multivariate effect size of rSVs $\hat{\boldsymbol{\gamma}}$ could be calculated by

$$
\begin{aligned}
\hat{\boldsymbol{\gamma}} &= \boldsymbol{D}^{-1}\hat{\boldsymbol{\beta}} \\
&= \boldsymbol{D}^{-1}(\boldsymbol{X}^T\boldsymbol{X})^{-1}\begin{pmatrix} \hat{\theta}_1\boldsymbol{x}_1^T\boldsymbol{x}_1 \\ \hat{\theta}_2\boldsymbol{x}_2^T\boldsymbol{x}_2 \\ \cdots \\ \hat{\theta}_P\boldsymbol{x}_P^T\boldsymbol{x}_P \end{pmatrix}.
\end{aligned} \tag{3}
$$

Because we also have

$$
\begin{aligned}
\hat{\boldsymbol{\gamma}} &= (\boldsymbol{T}^T\boldsymbol{T})^{-1}\boldsymbol{T}^T\boldsymbol{Y} \\
&= (\boldsymbol{T}^T\boldsymbol{T})^{-1}\begin{pmatrix} \boldsymbol{t}_1^T\boldsymbol{Y} \\ \boldsymbol{t}_2^T\boldsymbol{Y} \\ \cdots \\ \boldsymbol{t}_K^T\boldsymbol{Y}. \end{pmatrix}
\end{aligned} \tag{4}
$$

Therefore, by equation (3) and (4), we have

$$
\begin{pmatrix} \boldsymbol{t}_1^T\boldsymbol{Y} \\ \boldsymbol{t}_2^T\boldsymbol{Y} \\ \cdots \\ \boldsymbol{t}_K^T\boldsymbol{Y} \end{pmatrix} = \boldsymbol{T}^T\boldsymbol{T}\hat{\boldsymbol{\gamma}} \tag{5}
$$

Our final goal is to estimate the marginal effect size of rSV $\hat{\eta}_k, k = 1, ..., K$, which should be estimated in regression $y_i = t_{i,k}\eta_k + e_i$ and

$$\hat{\eta}_k = (\boldsymbol{t}_k^T \boldsymbol{t}_k)^{-1} \boldsymbol{t}_k^T \boldsymbol{Y} \tag{6}$$

Therefore, according to equation (6),

$$\begin{pmatrix} \hat{\eta}_1 \\ \hat{\eta}_2 \\ \dots \\ \hat{\eta}_K \end{pmatrix} = \begin{pmatrix} (\boldsymbol{t}_1^T \boldsymbol{t}_1)^{-1} \boldsymbol{t}_1^T \boldsymbol{Y} \\ (\boldsymbol{t}_2^T \boldsymbol{t}_2)^{-1} \boldsymbol{t}_2^T \boldsymbol{Y} \\ \dots \\ (\boldsymbol{t}_K^T \boldsymbol{t}_K)^{-1} \boldsymbol{t}_K^T \boldsymbol{Y} \end{pmatrix}. \tag{7}$$

When both $\boldsymbol{t}_1, ..., \boldsymbol{t}_K$ and $\boldsymbol{x}_1, ..., \boldsymbol{x}_P$ are standardized to mean 0 and variance 1, then

$$\begin{pmatrix} \hat{\eta}_1 \\ \hat{\eta}_2 \\ \dots \\ \hat{\eta}_K \end{pmatrix} = \frac{1}{N-1} \begin{pmatrix} \boldsymbol{t}_1^T \boldsymbol{Y} \\ \boldsymbol{t}_2^T \boldsymbol{Y} \\ \dots \\ \boldsymbol{t}_K^T \boldsymbol{Y} \end{pmatrix} = \frac{1}{N-1} \boldsymbol{T}^T \boldsymbol{T} \hat{\boldsymbol{\gamma}} = \boldsymbol{T}^T \boldsymbol{T} \boldsymbol{D}^{-1} (\boldsymbol{X}^T \boldsymbol{X})^{-1} \begin{pmatrix} \hat{\theta}_1 \\ \hat{\theta}_2 \\ \dots \\ \hat{\theta}_P \end{pmatrix} = \boldsymbol{D}^T \begin{pmatrix} \hat{\theta}_1 \\ \hat{\theta}_2 \\ \dots \\ \hat{\theta}_P \end{pmatrix}. \tag{8}$$
